# An electrostatic repulsion model of centromere organisation

**DOI:** 10.1101/2025.09.01.673455

**Authors:** Caelan Bell, Lifeng Chen, M. Julia Maristany, Claudia Blaukopf, Huabin Zhou, Jan Huertas, Jose I. Perez Lopez, Christoph C. H. Langer, Thomas L. Steinacker, Nikki Schütte, Lynda K Doolittle, Jorge R. Espinosa, Sy Redding, Rosana Collepardo-Guevara, Michael K. Rosen, Daniel W. Gerlich

**Affiliations:** Institute of Molecular Biotechnology of the Austrian Academy of Sciences (IMBA), Vienna BioCenter, 1030 Vienna, Austria; Vienna BioCenter PhD Program, Doctoral School of the University of Vienna and Medical University of Vienna, 1030 Vienna, Austria; Marine Biological Laboratory Chromatin Collaborative, Marine Biological Laboratory, Woods Hole, MA 02543, USA; Department of Biophysics and Howard Hughes Medical Institute, University of Texas Southwestern Medical Center, Dallas, TX 75390, USA; Maxwell Centre, Cavendish Laboratory, Department of Physics, University of Cambridge, Cambridge CB3 0HE, UK; Yusuf Hamied Department of Chemistry, University of Cambridge, Cambridge CB2 1EW, UK; Department of Physical-Chemistry, Universidad Complutense de Madrid, Av. Complutense s/n, 28040, Madrid, Spain; Department of Biochemistry and Molecular Biotechnology, University of Massachusetts Chan Medical School, Worcester, MA, USA; Department of Genetics, University of Cambridge, Cambridge CB2 3EH, UK

## Abstract

During cell division, chromosomes reorganise into compact bodies in which centromeres localise precisely at the chromatin surface^1–4^ to enable kinetochore-microtubule interactions essential for genome segregation^5–8^. The physical principles guiding this centromere positioning remain unknown. Here, we reveal that human core centromeres are directed to the chromatin surface by repulsion of centromere-associated proteins – independent of condensin-mediated loop extrusion and microtubule engagement. Using cellular perturbations, biochemical reconstitution, and multiscale molecular dynamics simulations, we show that chromatin surface localisation emerges from repulsion between condensed chromatin and both the kinetochore and the highly negatively charged centromere protein, CENP-B. Together, these elements form a centromeric region composed of two domains with opposing affinities, one favouring integration within the mitotic chromosome and the other favouring exposure to the surrounding cytoplasm, thereby driving surface positioning. Tethering synthetic negatively charged proteins to chromatin was sufficient to recapitulate this surface localisation in cells and *in vitro*, indicating that electrostatic repulsion is a key determinant of surface localisation. These findings demonstrate that centromere layering is not hardwired by chromatin folding patterns but instead emerges from phase separation in chromatin. Our work uncovers electrostatic polarity as a general and programmable mechanism to spatially organise chromatin.

---

The faithful segregation of chromosomes in dividing cells is essential for genome stability and inheritance. This process relies on the physical attachment of condensed chromatin to spindle microtubules. Spindle microtubules engage chromosomes at the centromere, a specialised chromosomal region enriched in the histone H3 variant CENP-A^9–13^. Core centromeric elements enriched in CENP-A are positioned at the surface of condensed mitotic chromatin^1–4^, where they serve as an assembly platform for kinetochores, large multiprotein complexes that physically link chromatin to the spindle^14,15^. Surface localisation of the centromere/kinetochore is crucial for efficient microtubule attachment, as spindle microtubules are unable to penetrate the dense mass of mitotic chromatin^16^. However, how chromatin fibres fold to position CENP-A-rich centromere cores at the chromosome surface is unknown. Elucidating this positioning mechanism is key to understanding how mitotic chromosome architecture fulfils the requirements for accurate genome segregation.

In human chromosomes, the CENP-A-rich centromere core typically spans ∼200–600 kilobases^11,17,18^, and it mediates kinetochore assembly via direct interactions with inner kinetochore components^4,9,10,12,15,19,20^. Upon mitotic entry, CENP-A-rich core domains relocate to the periphery of condensed chromatin, just beneath the kinetochore plate^1–4^. This layered centromere architecture has been proposed to arise from specific folding or looping of centromeric chromatin in a manner that brings CENP-A-rich domains to the chromosome surface^2,3,9,12,21,22^, yet the mechanisms underlying this organisation have not been definitively established.

Here, we examine how centromere organisation is governed by two fundamental processes that structure mitotic chromosomes: condensin-mediated DNA loop extrusion^23–26^ and chromatin phase separation^6,8,16,27–29^. We show that the positioning of the human core centromere at the mitotic chromosome surface arises through an electrostatic polarity mechanism, in which repulsion of kinetochore components and the centromere-associated CENP-B protein from dense chromatin promotes the phase boundary localisation of CENP-A-rich domains. This mechanism operates at least in part by electrostatic repulsion among negatively charged molecules and is independent of DNA looping. These data combined with *in vitro* chromatin engineering and multiscale molecular dynamics (MD) simulations reveal how intrinsic physicochemical properties of chromatin contribute to the three-dimensional architecture of centromeres.

## A condensin- and microtubule-independent mechanism targets centromeres to the chromatin surface

To precisely map CENP-A-rich domains relative to the bulk chromatin surface, we combined CENP-A immunofluorescence microscopy with semi-automated computational analysis (Fig. 1a,b; Fig. S1a,b). In prometaphase cells, this approach detected CENP-A foci essentially at the chromatin surface (average depth of 46 ± 107 nm (mean ± SD)), with only 5.7 ± 5.0% of foci located deeper than 200 nm (n = 450 foci across 30 cells) (Fig. 1a– d). These results confirm that centromeres adopt a layered architecture during mitosis, with CENP-A-rich domains confined to a narrow peripheral zone of the chromatin mass^1–4^.

**Fig. 1.**
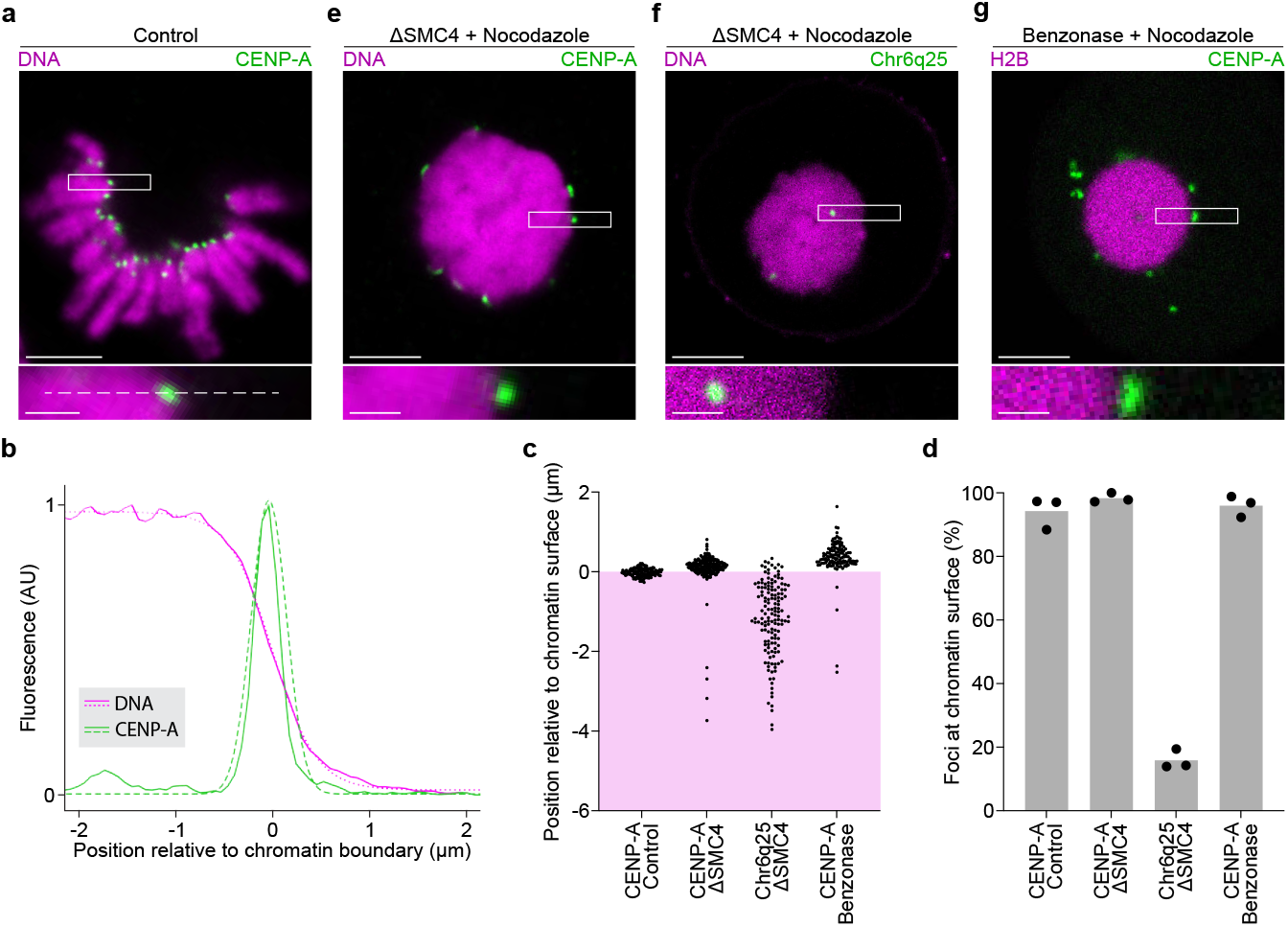
Centromere surface localisation is established independently of condensins and spindle microtubules. **a**, CENP-A localisation on unperturbed prometaphase chromosomes of HeLa cells, visualised by immunofluorescence and Hoechst-33342 staining. Inset showing line drawn across CENP-A focus and chromosome arm for quantification of CENP-A position relative to chromosome surface. **b**, Normalised fluorescence intensity profile for CENP-A and chromatin for line drawn in **a**, relative to chromatin surface position as demarcated by the half-maximum intensity of fitted chromatin curve at x=0. **c**, Quantification of CENP-A or FISH focus localisation relative to chromatin surface in cells treated as described in **a, e-g** from one biological replicate each. *n* = 135 (CENP-A control), *n* = 228 (CENP-A ΔSMC4), *n* = 144 (Chr6q25 ΔSMC4), *n* = 128 (CENP-A-EGFP, Benzonase). **d**, Quantification of percentage of foci localised at the chromatin surface as in **c** across 3 biological replicates. *n* = 135 / 150 / 165 (CENP-A control), *n* = 228 / 240 / 217 (CENP-A ΔSMC4), *n* = 144 / 36 / 35 (Chr6q25 ΔSMC4), *n* = 128 / 87 / 39 (CENP-A-EGFP, Benzonase), **e**, CENP-A localisation on mitotic chromatin mass after acute condensin-depletion (ΔSMC4) and microtubule depolymerisation by Nocodazole. Immunofluorescence image stained as in **a. f**, Localisation of Chr6q25 arm locus on ΔSMC4 and microtubule depolymerised mitotic chromatin mass, visualised by RASER-FISH; DNA was stained by DAPI. **g**, Localisation of CENP-A on liquid chromatin condensate, induced by Benzonase microinjection in Nocodazole-arrested mitotic cell. Live-cell image with CENP-A visualised by EGFP-tag; Chromatin was visualised by H2B-mCherry. Scale bars: 5 µm, 1µm for insets (**a, e**-**g**). *n* indicates number of foci per biological replicate of 10 cells (for Chr6q25, per biological replicate of 39 / 10 / 10 cells).

Condensins are key regulators of mitotic chromosome architecture^6–8,25,26^, providing mechanical stability^30–33^ and organising core centromeres into a bipartite structure on each sister chromatid^34^. Whether condensins also contribute to the surface targeting of centromeres has remained unclear. To address this, we depleted the SMC4 subunit, which is shared by both condensin I and II complexes, using a homozygous degron-tagged cell line^16^. SMC4 degradation was induced in G2-synchronised cells, followed by release into mitosis in the presence of Nocodazole to depolymerise microtubules and prevent spindle-mediated distortions of chromosomes (Fig. S1c-f).

Under these conditions, chromatin formed a single compact mass, whereby the mitotic state was verified by immunofluorescence detection of Ser10 phosphorylation on histone H3, while cleaved-caspase staining confirmed that the cells had not undergone apoptosis (Fig. S1g-i). CENP-A foci remained predominantly localised at the chromatin surface even in the absence of condensins: 98.4% ± 1.5% of foci were within 200 nm of the surface (Fig. 1c–e). In contrast, fluorescence in situ hybridisation (FISH) to two chromosome arm loci (Chr6q25 and Chr8q22.2) revealed that only 15.9% ± 3.1% and 6.9% ± 6.7% of signals, respectively, were similarly positioned near the chromatin periphery, with many located several µm deep within the interior (Fig. 1c,d,f; Fig. S1j-l). These results demonstrate that CENP-A-rich centromere cores are specifically targeted to the chromatin surface through a mechanism that operates independently of condensins and spindle forces.

This condensin independence suggests that surface localisation arises from mechanisms unrelated to long-range chromatin folding. We hypothesised that surface targeting might result from the intrinsic ability of chromatin to compact and phase separate, both *in vitro* and in mitotic cells^16,27,28^. Notably, condensation of mitotic chromatin is preserved even when chromatin fibres are fragmented by injecting endonuclease into mitotic cells, which transforms chromosomes into spherical droplets^16^. This procedure provides an assay system to explore the role of chromatin phase separation in surface targeting – independent of long-range loop architectures.

To test whether CENP-A domains remain surface-localised in the absence of intact chromatin fibres, we microinjected the endonuclease Benzonase into Nocodazole-treated mitotic cells. As previously observed^16^, this treatment converted rod-shaped chromosomes into spherical chromatin condensates (Fig. S2a,b). Fluorescence recovery after photobleaching (FRAP) of H2B-mCherry confirmed that the chromatin condensates were liquid (Fig. S2c,d). Live-cell imaging of GFP-tagged CENP-A revealed that 96.0 ± 3.4% of CENP-A foci remained localised at the condensate surface (Fig. 1c,d,g). The layered organisation of centromeres thus does not depend on the integrity of chromatin fibres nor on engagement of kinetochores with microtubules.

Together, these findings demonstrate that the surface localisation of centromeres does not require condensin-mediated chromosome folding or long-range chromatin loop structures. Instead, they suggest that CENP-A-rich domains are directed to the chromatin surface via physical properties associated with condensed chromatin. This challenges prior models proposing loop- or helix-based centromere positioning^2,3^, and supports an alternative framework in which centromere layering arises from local interactions in condensed chromatin.

### Centromere layering depends on kinetochores and CENP-B

The positioning of CENP-A domains might be influenced by the associated kinetochore proteins. To investigate this possibility, we depleted CENP-C, a core component of the constitutive centromere-associated network (CCAN) that is essential for recruitment of all other kinetochore subunits^20,35,36^. CENP-C depletion was achieved by combining Cas9 ribonucleoprotein complexes and siRNAs, alongside microtubule depolymerisation and condensin depletion as before (Fig. S3a,b). Efficient kinetochore disruption was confirmed by immunofluorescence using NDC80 and KNL1 as a markers for the outer kinetochore (Fig. 2a,b; Fig. S3c,d). Quantification of CENP-A foci positions revealed a pronounced redistribution of centromeres following kinetochore depletion (Fig. 2a,c,d). In control cells, 98.5% ± 1.5% of CENP-A foci localised within 200 nm of the chromatin surface, whereas upon CENP-C depletion, only 40.6% ± 6.6% remained surface-associated, indicating that kinetochores are critical for targeting CENP-A-rich chromatin to the phase boundary of condensed chromatin.

**Fig. 2.**
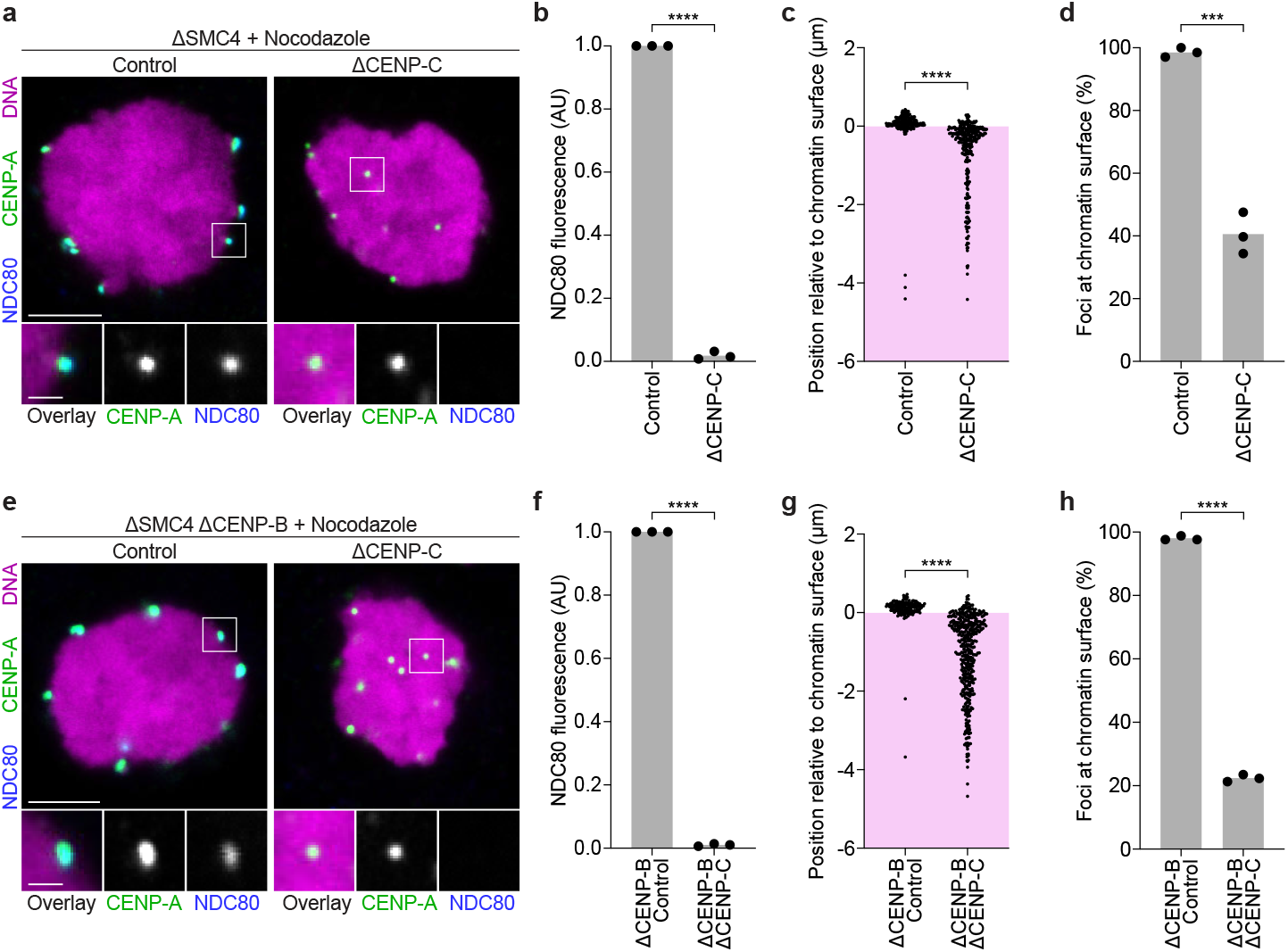
Kinetochore complexes and CENP-B cooperatively establish centromere surface localisation. **a**, Localisation of CENP-A foci upon kinetochore depletion (ΔCENP-C) in condensin-depleted (ΔSMC4), microtubule depolymerised cells arrested in mitosis by proTAME. NDC80 signal was used as readout of kinetochore depletion. **b**, Quantification of kinetochore depletion by mean NDC80 fluorescence at CENP-A foci used in localisation quantification (**c**) as in **a** per biological replicate (*n* = 206 / 244 / 251 (control), *n* = 284 / 269 / 200 (ΔCENP-C)). Significance was tested using a two-tailed t-test (*P* < 10^-15^ (precision limit of floating-point arithmetic), t(4) = 138.8). **c**, Quantification of CENP-A foci localisation relative to chromatin surface for ΔCENP-C and control cells as in **a** from one biological replicate (*n* = 206 (control), *n* = 284 (ΔCENP-C)). Significance was tested using a two-tailed Mann-Whitney *U*-test (*P* < 10^-15^ (precision limit of floating-point arithmetic), U = 5728). **d**, Quantification of percentage of CENP-A foci localised at the chromatin surface across biological replicates (*n* = as in **b**). Significance was tested using a two-tailed t-test (*P* = 0.000121, t(4) = 14.8). **e-h** as in **a-d** but in ΔCENP-B cells. *n* = 180 / 217 / 210 (ΔCENP-B control), *n* = 430 / 378 / 310 (ΔCENP-B ΔCENP-C). **f**, Significance was tested using a two-tailed t-test (*P* < 10^-15^ (precision limit of floating-point arithmetic), t(4) = 438.4). **g**, Significance was tested using a two-tailed Mann-Whitney *U*-test (*P* < 10^-15^ (precision limit of floating-point arithmetic), U = 4369). **h**, Significance was tested using a two-tailed t-test (*P* 10^-15^ (precision limit of floating-point arithmetic), t(4) = 98.4). Images are linearly contrast adjusted and matched between conditions. Scale bars: 5 µm, 1 µm for insets (**a, e**). *n* indicates number of foci per biological replicate of 10 cells.

Although CENP-C depletion markedly reduced surface localisation, a subset of CENP-A domains remained at the chromatin surface in the absence of kinetochores. We hypothesised that this residual localisation could be mediated by centromere proteins that function independently of CENP-C. One such candidate is CENP-B, a DNA-binding protein that recognises CENP-B-box motifs within alpha-satellite DNA^37,38^. While these motifs are also present in pericentromeric regions flanking the CENP-A–rich core centromere, DNA methylation in pericentromeric chromatin suppresses CENP-B binding in these domains, resulting in enrichment of CENP-B at the core centromere^39–41^. To assess its potential contribution, we generated CENP-B knockout cells in an SMC4-degron background (Fig. S3e,f). CENP-B knockout alone had no detectable effect on CENP-A domain localisation; however, when combined with kinetochore-depletion, surface localisation was further reduced, with only 22.4% ± 1.1% of CENP-A foci remaining at the surface (Fig. 2e-h). These findings demonstrate that centromere layering is cooperatively established by kinetochore complexes and CENP-B, suggesting a model in which CENP-A-rich chromatin is brought to the chromatin surface by its associated proteins. While kinetochores play a dominant role in positioning CENP-A chromatin at the condensate surface, CENP-B provides a parallel mechanism that partially compensates when kinetochores are absent.

### CENP-B promotes chromatin surface localisation in vitro via its acidic domain

To determine whether centromere-associated proteins are sufficient to mediate surface targeting and to understand the underlying physical principles, we established a reconstituted *in vitro* assay based on phase-separated chromatin and purified recombinant candidate proteins. We used dodecameric nucleosome arrays that undergo liquid–liquid phase separation under physiological salt conditions, forming condensates that recapitulate condensin-independent chromatin compaction observed during mitosis^16,27,28^. We first examined whether substituting histone H3 with CENP-A alters the spatial localisation of nucleosome arrays within chromatin condensates. CENP-A-based nucleosome arrays mixed homogeneously with those based on the canonical histone H3, indicating that CENP-A alone is insufficient to produce layered chromatin condensate structures (Fig. S4a,b). This finding is consistent with the cellular requirement of kinetochores and CENP-B for centromeric surface localisation.

To investigate the potential roles of other centromere-associated proteins, we engineered nucleosome arrays containing defined protein-binding sites. Specifically, to assess the role of CENP-B, we designed nucleosome arrays with an appended 109-base pair DNA sequence containing four CENP-B-boxes. When mixed at a 1:10 ratio with unmodified canonical nucleosome arrays, these CENP-B-box-containing arrays were uniformly distributed throughout the condensates (Fig. 3a,b). In the presence of CENP-B, however, the CENP-B-box-containing arrays were depleted from the interior and relatively enriched at the condensate periphery (Fig. 3a,b). Addition of excess competitor DNA containing CENP-B-boxes displaced CENP-B from the surface and unbound CENP-B-box-containing chromatin arrays localised inside the condensate (Fig. S5a), confirming that CENP-B binding to the target array is necessary for chromatin surface targeting. Together, these results demonstrate that CENP-B is sufficient to drive its bound nucleosome arrays to the periphery of chromatin condensates.

**Fig. 3.**
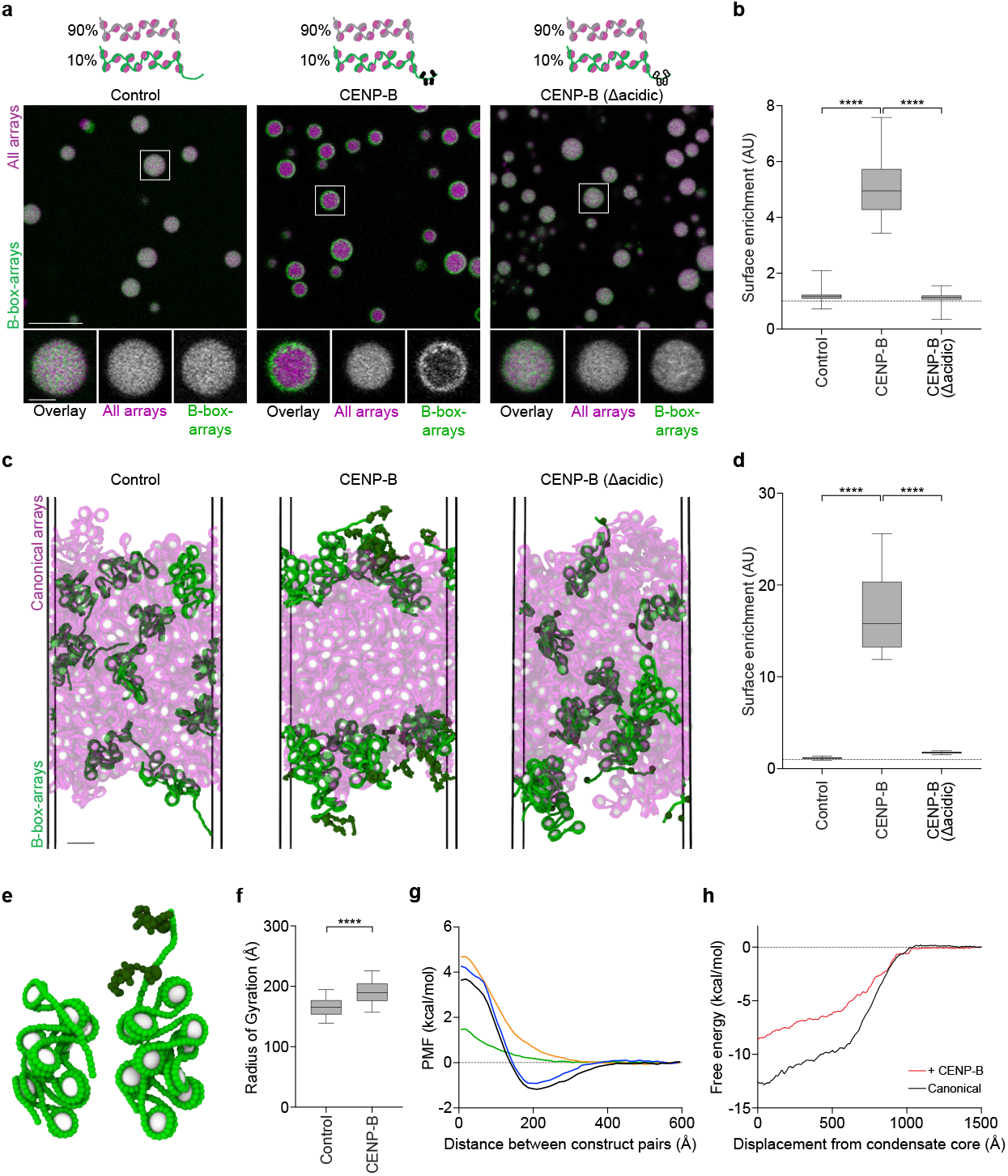
CENP-B acidic patch drives surface localisation of bound chromatin. **a**, Chromatin condensates of canonical nucleosome arrays mixed at 10:1 ratio with CENP-B-box-containing arrays (B-box-arrays), without protein (control), with full-length CENP-B or with CENP-B lacking the acidic region (Δacidic). All arrays stained with Hoechst-33342, B-box-arrays additionally visualised by 5’ Cy5. Scale bars: 10 µm, 2 µm for inset. **b**, Quantification of B-box-arrays surface enrichment for condensates as in **a**. *n* = 395 (Control), *n* = 744 (CENP-B), *n* = 674 (CENP-B (Δacidic)); *n* indicates number of condensates per condition. Significance was tested using a two-tailed t-test (Control vs CENP-B: *P* < 10^-15^ (precision limit of floating-point arithmetic), t(1137) = 64.7; CENP-B vs CENP-B (Δacidic): *P* < 10^-15^ (precision limit of floating-point arithmetic), t(1416) = 85.3). **c**, Representative configuration of coarse-grained chromatin condensates obtained from Direct Coexistence MD simulations in a rectangular 3D box with one long axis, where condensed and dilute phases coexist separated by an interface. Shown are canonical chromatin arrays and B-box-arrays under the conditions in **a**. Scale bar: 200 Å. **d**, Quantification of B-box-array surface enrichment for simulations in **c**, averaged over 20 independent frames per condition. Significance was tested using a two-tailed t-test (Control vs CENP-B: *P* < 10^-15^ (precision limit of floating-point arithmetic, t(38) = 17.6; CENP-B vs CENP-B (Δacidic): *P* < 10^-15^ (precision limit of floating-point arithmetic), t(38) = 16.9). **e**, Representative configurations of coarse-grained nucleosome arrays showing unbound B-box-arrays (left) and CENP-B-bound arrays (right). CENP-B is shown in dark green. **f**, Radius of gyration for chromatin arrays with and without CENP-B. See methods for details on *n*. Significance was tested using a two-tailed t-test (Control vs CENP-B: *P* < 10^-15^ (precision limit of floating-point arithmetic, t(1002) = 20.6). **g**, Free-energy change associated with forming interactions between pairs of different portions of the fibre constructs, represented by PMFs computed from umbrella sampling MD simulations. The intermolecular centre-of-mass distance was used as the reaction coordinate, and PMFs were referenced to zero at large separations. Shown are values for pairs of canonical nucleosome-associated regions (black), pairs of canonical nucleosome-associated regions and CENP-B-bound DNA-fragments (green), pairs of bound DNA-fragments (orange) and pairs of canonical nucleosome-associated regions ligated to bound DNA-fragments (blue). **h**, Free-energy change associated with extracting either a canonical array (black) or a CENP-B–bound array (red) from the condensate centre. The centre-of-mass position of the extracted array measured from the centre of the condensate was used as the reaction coordinate, and PMF were referenced to zero at large distances. Magenta shading denotes the condensate interior.

To elucidate the mechanism of these CENP-B– mediated effects, we dissected the domains necessary for function. The dimerisation domain of the protein is important for centromere structure and compaction^42^. To test whether dimerisation is required for surface targeting, we generated a truncated version of CENP-B lacking the dimerisation domain (CENP-B (Δdimer); Fig. S5b). This mutant recapitulated the effects of wild-type CENP-B: target arrays were excluded from the condensate interior and relatively enriched at the surface (Fig. S5c,d), indicating that CENP-B-dimerisation is not needed for surface targeting.

CENP-B contains one of the longest acidic stretches in the human proteome (Fig. S5b). Previous studies have shown that negatively charged proteins are typically excluded from condensed chromatin both *in vitro* and in cells^16,43^, suggesting a potential role of electrostatic interactions in surface targeting. To investigate this, we generated a truncated version of CENP-B lacking the acidic region (CENP-B (Δacidic); Fig. S5b) and assessed its ability to promote surface targeting in our *in vitro* reconstitution assay. Unlike the full-length protein, in the presence of the Δacidic mutant, CENP-B-box-containing arrays remain homogeneously distributed within the condensate interior (Fig. 3a,b). These findings demonstrate that the acidic domain is essential for CENP-B-mediated chromatin surface localisation.

To determine the physical basis of CENP-B-mediated chromatin surface targeting, we performed MD simulations of chromatin condensates. These simulations employ a multiscale framework that bridges atomistic detail with mesoscale organisation to derive a minimal chromatin model^44^, which accurately reproduces the stability and material properties of chromatin condensates^45^ and captures the structure of chromatin arrays within them^46^. To represent CENP-B, we derived a consistent minimal model from its atomistic structure and enforced its binding to B-box DNA sequences (see Methods). As in the experiments, we simulated mixtures of unmodified nucleosome arrays with arrays containing extended DNA regions with B-box sequences. In the absence of CENP-B, chromatin arrays with or without B-boxes remained well mixed within simulated condensates (Fig. 3c,d). By contrast, simulations including wild-type CENP-B bound to chromatin showed a clear enrichment of CENP-B-bound arrays at the condensate surface. This surface enrichment was abolished in simulations with CENP-B lacking the acidic domain, consistent with experimental observations (Fig. 3c,d). Our multiscale framework thus accurately recapitulates the CENP-B-induced surface targeting of associated chromatin, providing a powerful tool for further dissection of the underlying mechanism.

To investigate how CENP-B influences the structure of individual chromatin arrays, we analysed their configurational ensembles from simulated nucleosome arrays. In the absence of CENP-B, the DNA segment containing B-box sequences folded back onto the nucleosome array, driven by associative interactions with the positively charged histone core (Fig. 3e,f). In contrast, binding of full-length CENP-B induced an extended conformation, with both the B-box DNA and the bound CENP-B protruding away from the array (Fig. 3e,f). These findings suggest that CENP-B binding imposes structural polarity, orienting the B-box DNA and the bound CENP-B away from the condensate interior. Notably, unlike nucleosomal DNA, whose negative charge is partially neutralised by the positively charged histone proteins, the extended B-box-CENP-B region carries a higher net negative charge, potentially enhancing electrostatic repulsion from the chromatin-dense condensate core. Together, these features create a local charge asymmetry along the chromatin fibre, which can drive directional organisation within condensates.

To mechanistically understand CENP-B-bound chromatin surface enrichment, we mapped the free-energy landscape of interactions between different portions of the fibre construct by calculating the potential of mean force (PMF) as a function of their separation (Fig. 3g). Interactions between two canonical nucleosome-associated regions were attractive, as indicated by the pronounced negative dip in the PMF. By contrast, any interaction that involved the CENP-B-bound DNA segment was repulsive: both the canonical array with the CENP-B-bound DNA fragment and the CENP-B-bound DNA fragment with another bound fragment yielded positive PMF values across all separations examined. A construct that contained both a nucleosome-associated region plus a CENP-B-bound segment retained attraction but with reduced magnitude relative to canonical nucleosome-only pairs, consistent with an overall weakening of cohesive forces. Together, these PMF profiles indicate an anisotropic, polar architecture of CENP-B-bound chromatin, in which one end of the fibre is repulsive while the other is attractive, such that the lowest free-energy configuration is achieved when the repulsive CENP-B-bound end is exposed at the condensate interface, thereby leading to surface enrichment.

To further investigate why CENP-B-bound chromatin arrays preferentially partition to the condensate surface, we computed PMF profiles quantifying the change in free energy associated with extracting canonical versus CENP-B-bound arrays from the condensate. The free-energy cost of extraction was higher for canonical arrays (ΔG = 8.4) than for CENP-B-bound arrays (ΔG = 5.9; Fig. 3h), indicating that incorporating CENP-B-bound arrays deep into the condensate interior would decrease its thermodynamic stability. This finding, together with the pairwise PMFs (Fig. 3g), reveals the thermodynamic basis for CENP-B-mediated surface localisation. By localising at the condensate surface, CENP-B-bound arrays play a dual role in lowering the overall free energy of the condensate and enhancing its thermodynamic stability: (i) permitting the condensate interior to be preferentially occupied by canonical arrays, which engage in stronger attractive interactions and maximise the overall enthalpic gain of the condensate, and (ii) reducing the free-energy cost of interface formation via the enrichment of the repulsive CENP-B-bound DNA segments at the interface.

Together, our biochemical, cellular, and computational data converge on a model in which CENP-B drives the enrichment of centromeric chromatin at the chromatin– cytoplasm interface through charge-based repulsion from condensed chromatin. By tethering a highly acidic domain to chromatin, CENP-B introduces local electrostatic polarity, with one region favouring chromatin binding and an adjacent region disfavouring binding. We use the term electrostatic polarity to denote a directional asymmetry in the charge distribution of chromatin, distinct from molecular dipoles or electronic polarisation, whereby one end of the chromatin carries a higher net negative charge than the rest. Although the multi-subunit complexity of kinetochores currently limits reconstitution or dynamic simulations, *in silico* charge estimates based on amino acid composition at physiological pH predict an average net charge of -303.9e per kinetochore complex (Table S1), supporting a broader role for electrostatic polarity in organising centromeric chromatin layers during mitosis.

### Synthetic negative charge targets chromatin to the surface

As a principle grounded in fundamental physics, electrostatic repulsion may act as a more general mechanism of chromatin organisation - one that operates independently of specific protein structures or evolutionarily conserved molecular interactions. To test whether charge asymmetry alone is sufficient to induce chromatin surface targeting, we explored alternative strategies to locally increase negative charge. Double-stranded DNA carries a net charge of approximately –2e per base pair, suggesting that extended stretches of non-nucleosomal DNA could contribute substantial negative charge to impose electrostatic polarity on the chromatin fibres. To investigate this, we performed MD simulations in which nucleosome arrays were fused to nucleosome-free DNA segments of varying lengths and assessed their localisation relative to condensates formed by canonical nucleosome arrays. The simulations showed that free DNA segments of 300 bp consistently directed the associated nucleosome arrays to the condensate surface, with an even stronger effect observed for 500 bp segments. In contrast, 100 bp of free DNA was insufficient to induce surface targeting (Fig. 4a,b). Notably, the free DNA extended into the surrounding buffer, adopting an elongated conformation reminiscent of the orientation and repulsion observed for CENP-B bound to shorter DNA segments (Fig. S6a,b). Together, these findings suggest that negatively charged DNA extensions are sufficient to drive surface targeting of chromatin condensates through electrostatic exclusion.

**Fig. 4.**
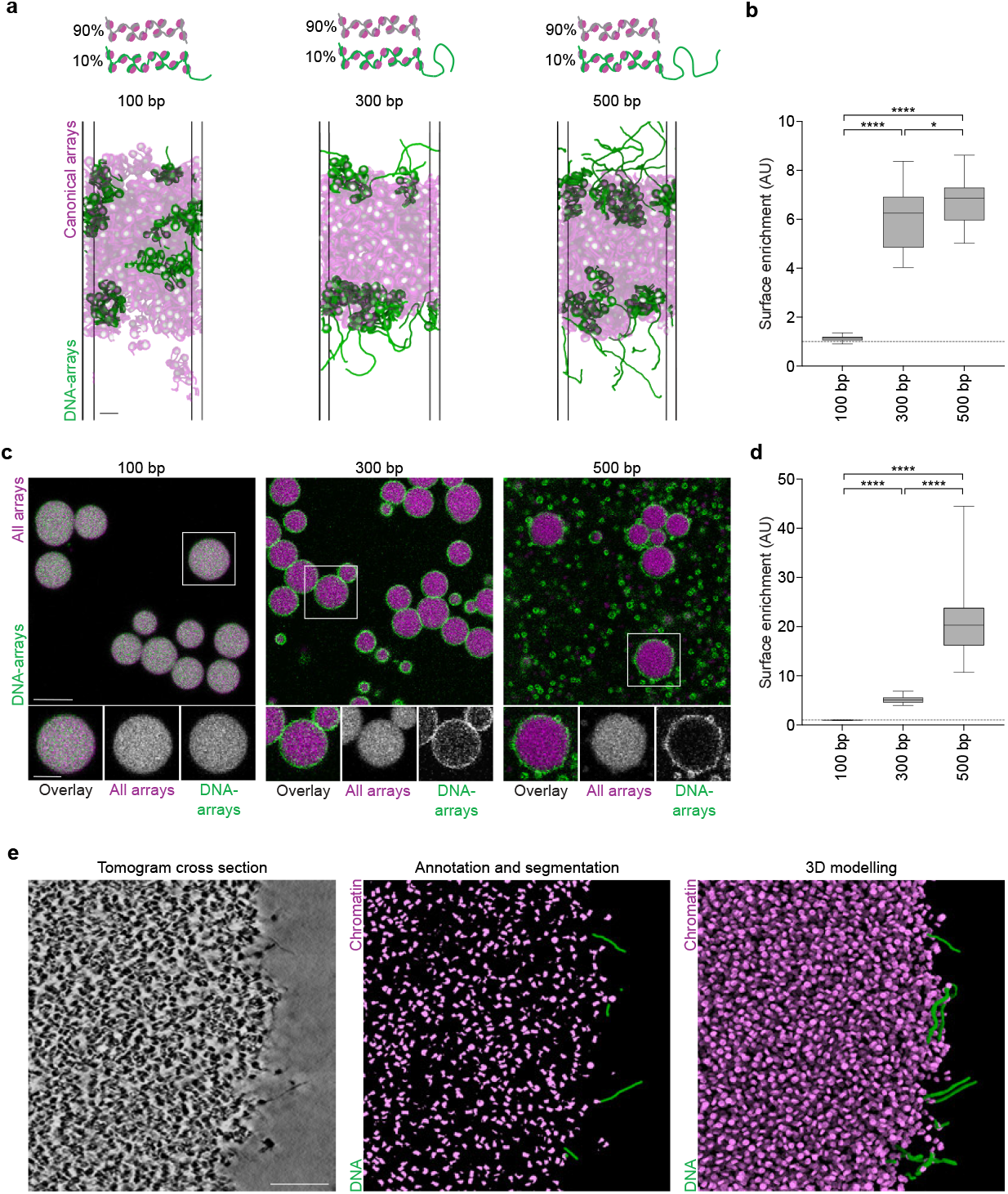
Electrostatic repulsion and polarity induce chromatin surface targeting. **a**, Representative configurations of coarse-grained chromatin condensates obtained from Direct Coexistence MD simulations in a rectangular 3D box with one long axis, containing 90% canonical nucleosome arrays and 10% arrays fused to nucleosome-free DNA segments of varying lengths (DNA-arrays). Scale bar: 200 Å. **b**, Quantification of DNA-arrays surface enrichment for simulations as in **a** across 20 snapshots per condition. Significance was tested using a two-tailed t-test (100 bp vs 300 bp: *P* < 10^-15^ (precision limit of floating-point arithmetic), t(38) = 17.4; 300 bp vs 500 bp: *P* = 0.0292, t(38) = 2.3; 100 bp vs 500 bp: *P* < 10^-15^ (precision limit of floating-point arithmetic), t(38) = 26.4). **c**, Chromatin condensates with experimental conditions as in **a**, All arrays stained with Hoechst-33342, DNA-arrays additionally visualised by 5’ Cy5. Scale bars: 10 µm, 2 µm for inset. **d**, Quantification of DNA-arrays surface enrichment for condensates as in **a**. *n* = 783 (100 bp), *n* = 560 (300 bp), *n* = 54 (500 bp); *n* indicates number of condensates per condition. Significance was tested using a two-tailed t-test (100 bp vs 300 bp: *P* < 10^-15^ (precision limit of floating-point arithmetic), t(1341) = 127.7; 300 bp vs 500 bp: *P* < 10^-15^ (precision limit of floating-point arithmetic), t(612) = 40.1; 100 bp vs 500 bp: *P* < 10^-15^ (precision limit of floating-point arithmetic), t(835) = 62.5). **e**, Representative cryo-ET structure of a chromatin condensate of nucleosome arrays ligated to 300 bp DNA, as in **c**. From left to right: representative tomographic slice (0.8 nm thick), corresponding section annotated with nucleosomes detected by context-aware template matching and manually segmented DNA, and resulting 3D model showing nucleosomes and traced DNA within the tomogram. Scale bar: 100 nm.

In parallel with the simulations we generated nucleosome arrays fused to negatively charged double-stranded DNA segments. As observed in the simulations, dodecameric arrays fused to 300 bp of naked DNA prominently accumulated at the condensate surface, whereas arrays fused to 100 bp of DNA were uniformly distributed throughout the condensates (Fig. 4c,d). Increasing the DNA length to 500 bp further enhanced surface accumulation, resulting in greater exclusion from the condensate interior (Fig. 4c,d). These findings support a charge-based repulsion mechanism that functions independently of specific protein–protein interactions. Moreover, they demonstrate that the extent of surface targeting can be modulated by tuning the negative charge, suggesting a potential regulatory mechanism in chromatin organisation.

The simulations also predicted that nucleosome arrays ligated to free DNA adopt an extended molecular conformation and orient consistently relative to the condensate surface. To test this, we performed cryo-electron tomography (cryoET) on arrays containing 300 bp of free DNA. The resulting images confirmed the presence of extended DNA elements protruding from the condensate surface (Fig. 4e), providing additional support for a model in which surface targeting arises from a balance between attractive and repulsive electrostatic forces.

To further probe the generic role of electrostatic polarity in chromatin surface targeting, we designed a synthetic protein-based construct using components with no homology to human proteins, thereby minimising the potential effect of stereospecific protein-protein interactions. The construct fused two functional domains: (1) the bacterial tetracycline repressor (TetR), which binds tetracycline operator (TetO) sequences with high affinity and specificity both *in vitro* and *in vivo*^47^, and (2) an engineered variant of GFP with an overall net charge of – 25 e at physiological pH, GFP(neg25)^48^. These domains were joined by a flexible linker to create a synthetic chromatin surface-targeting construct: TetR-GFP(neg25).

We first assessed whether this construct could promote chromatin surface enrichment *in silico*. Simulations were performed using nucleosome dodecamers carrying three TetO motifs, modelled with atomic-resolution input structures and coarse-grained dynamics. In control simulations, TetO-containing arrays mixed with canonical arrays remained homogeneously distributed throughout the condensate (Fig. 5a,b). Upon binding of TetR-GFP(neg25), however, TetO-arrays became enriched at the condensate surface, indicating that the addition of local negative charge biases associated chromatin toward the phase boundary. Notably, simulations in which TetR-GFP(neg25) was present but unbound did not result in surface accumulation (Fig. 5a,b), confirming that chromatin tethering is required for this effect.

**Fig. 5.**
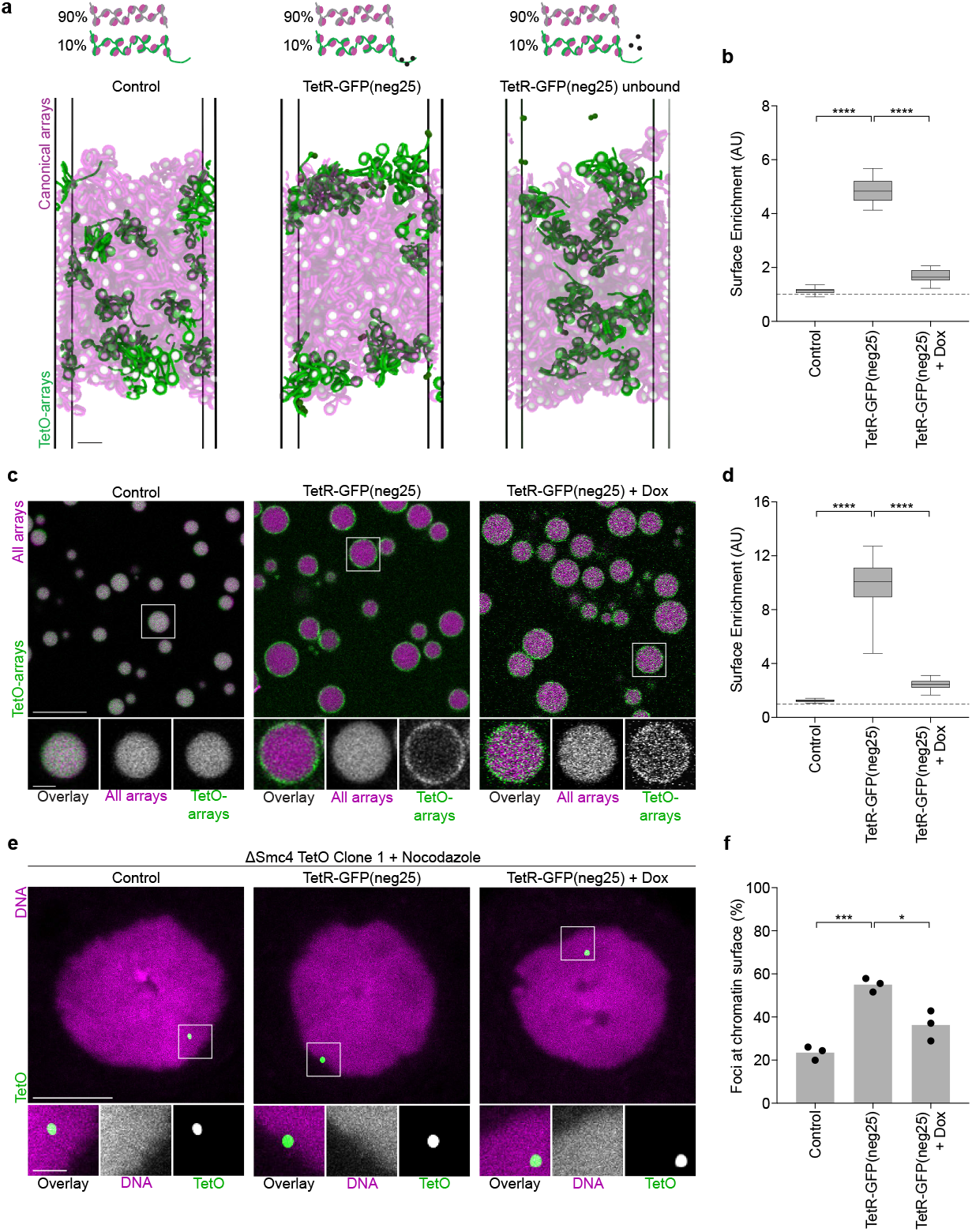
Electrostatic polarity through protein binding promotes chromatin surface enrichment. **a**, Representative configurations of coarse-grained chromatin condensates obtained from Direct Coexistence MD simulations in a rectangular 3D box with one long axis, containing 90% canonical nucleosome arrays and 10% TetO-containing nucleosome arrays (TetO-arrays), shown under three conditions: in the absence of protein (control), with TetR-GFP(neg25) bound to TetO-arrays or in the presence of unbound TetR-GFP(neg25). Scale bar: 200 Å. **b**, Quantification of TetO-array surface enrichment from simulations in **a**. Significance was tested using a two-tailed t-test (Control vs TetR-GFP(neg25): *P* < 10^-15^ (precision limit of floating-point arithmetic), t(38) = 36.6; TetR-GFP(neg25) vs TetR-GFP(neg25) unbound: *P* < 10^-15^ (precision limit of floating-point arithmetic), t(38) = 27.8). **c**, Chromatin condensates with experimental conditions as in **a**, using Doxycycline (Dox) to suppress binding of TetR-GFP(neg25) to TetO. All arrays stained with Hoechst-33342, TetO-arrays additionally visualised by 5’ Cy5. Scale bars: 10 µm, 2 µm for inset. **d**, Quantification of TetO-array surface enrichment for condensates as in **a**. *n* = 176 (control), *n* = 119 (TetR-GFP(neg25)), *n* = 282 (TetR-GFP(neg25) + Dox); *n* indicates number of condensates per condition. Significance was tested using a two-tailed t-test (Control vs TetR-GFP(neg25): *P* < 10^-15^ (precision limit of floating-point arithmetic), t(293) = 61.6; TetR-GFP(neg25) vs TetR-GFP(neg25) + Dox: *P* < 10^-15^ (precision limit of floating-point arithmetic), t(399) = 42). **e**, Localisation of TetO-repeats integrated into the genome of HeLa cells. The images show mitotic cells with depolymerised microtubules and ΔSMC4, with TetO visualised by RASER-FISH and DNA by DAPI. Conditions shown are: no TetR-GFP(neg25) protein (control), TetO bound by TetR-GFP(neg25), and presence of unbound TetR-GFP(neg25) + Dox. Scale bars: 5 µm, 1 µm for inset. **f**, Quantification of percentage of foci localised at the chromatin surface as in **e** per biological replicate across conditions. *n* = 45 / 30 / 23 (control), *n* = 72 / 31 / 19 (TetR-GFP(neg25)), *n* = 32 / 38 / 28 (TetR-GFP(neg25) + Dox), with *n* indicating number of foci/cells per biological replicate. Significance was tested using a two-tailed t-test (Control vs TetR-GFP(neg25): *P* = 0.003, t(4) = 12.2; TetR-GFP(neg25) vs TetR-GFP(neg25) + Dox: *P* = 0.0134, t(4) = 4.2).

To experimentally validate these predictions, we assembled TetO-containing nucleosome arrays and mixed them with canonical nucleosome arrays in phase separation assays. In the absence of TetR-GFP(neg25), TetO-arrays were uniformly distributed within the condensate interior. In the presence of TetR-GFP(neg25), TetO-arrays were depleted from the condensate interior and were relatively surface enriched (Fig. 5c,d). This effect was reversed by the addition of doxycycline, which disrupts TetR-TetO binding, confirming that surface targeting depends on specific chromatin tethering (Fig. S7a).

To investigate how tethering charge influences chromatin positioning in cells, we inserted approximately 200 TetO-repeats at a random genomic location in SMC4-degron HeLa cells. In mitotic cells depleted of condensins and spindles, FISH showed that the TetO-repeat localised predominantly to the chromatin interior (Fig. 5e,f), consistent with the typical positioning of chromosome arms (see Fig. 1). Upon expression of TetR-GFP(neg25), a significantly higher proportion of TetO FISH foci localised to the chromatin periphery (Fig. 5e,f, Fig. S7b,c). This surface enrichment was reduced by doxycycline treatment (Fig. 5e,f, Fig. S7b,c). Similar results were obtained across two additional independent TetO-integrant clones (Fig. S7d-g). These observations demonstrate that surface targeting requires neither an evolved chromatin ligand nor a specific genomic locus. While these findings qualitatively mirror the *in vitro* experiments, the extent of surface enrichment was less pronounced in cells. This attenuation may reflect the more stringent constraints on chromatin diffusion imposed by the continuity of the chromatin fibre in intact chromosomes.

Together, these findings demonstrate that local tethering of negative charge is sufficient to drive chromatin surface localisation in both reconstituted systems and living cells. The ability of a completely synthetic system to recapitulate this behaviour underscores the generality of electrostatic polarity as a principle of chromatin organisation.

## Conclusions

Our study uncovers a physical mechanism by which the layered architecture of human centromeres is established during mitosis. In contrast to prevailing models that attribute centromere positioning to specific chromatin loop configurations^2,3,9,12,21,22^, we demonstrate that the surface localisation of CENP-A-rich domains is driven by the balance between intrinsic chromatin-chromatin attraction within mitotic chromosomes and repulsive forces generated by kinetochore complexes and CENP-B. These opposing forces are, at least in part, governed by local electrostatic interactions, which originate from the negatively charged chromatin fibre^27–29^ and negatively charged proteins tightly bound to chromatin (Fig. 5). This form of interfacial targeting through intrinsic electrostatic polarity—that is, a directional charge asymmetry along chromatin— is conceptually analogous to the behaviour of amphiphilic molecules that localise at the interface between aqueous and lipid phases^49^.

Our ability to recapitulate surface targeting using a synthetic, highly charged fusion protein demonstrates that electrostatic polarity is a modular and programmable feature of chromatin that can be engineered independently of endogenous centromere components. Notably, the observation that tethering acidic peptides can relocate chromosomal loci from the nuclear periphery to the interior during interphase, when chromatin is not homogeneously compacted as in mitosis^50^ suggests that electrostatic polarity may serve as a general mechanism governing chromosome architecture beyond mitosis.

Together, these findings redefine our understanding of centromere architecture and open new avenues for exploring how physical forces and charge asymmetries govern genome organisation. Future studies will be essential to elucidate how these physical principles integrate with chromosome organisation by loop extrusion, cohesion, and kinetochore clustering. It will be important to investigate whether similar layering processes operate on chromatin during interphase, and how dynamic regulation of chromatin charge contributes to genome function. More broadly, the capacity to engineer chromatin positioning through electrostatic design offers promising avenues for synthetic genome control and compartment-specific epigenetic regulation. Overall, our results underscore the emerging importance of affinity-based and phase separation–driven mechanisms in shaping chromatin architecture.

## Acknowledgements

We thank Daniel and David Isenberg for their generous support of the MBL Chromatin Collaborative in honor of their father, Irvin Isenberg. We thank I. Patten and M. F. D. Spicer for comments on the manuscript. We further thank J.N. Cruz and the IMBA/ IMP/GMI BioOptics and Molecular Biology Service facilities for technical support, J. Ellenberg and K.S. Beckwith for technical advice on RASER-FISH, and M.W.G. Schneider, S. Kolesnikova, and T. Nagaraju for preliminary observations. We thank I.M. Cheeseman for providing antibodies against kinetochore proteins. We thank A. Lyon for assistance with *in vitro* image analyses.

## Author contributions

CBell, LC, MJM, SR, RCG, MKR, and DWG conceived the study. CBell performed cell biological experiments, except TetO/TetR experiments (by CBlaukopf). LC performed biochemical experiments, except cryoET experiment (by HZ). MJM, JH, JIPL, JRE performed modelling and computer simulations. CBell, LC, LD, NS, and TLS designed and generated reagents. SR developed and applied scripts for biochemical image analysis. CCHL developed strategy for cell biological image analysis and developed corresponding software, CBell created jupyter notebooks applying the software for image analysis in cells. DWG and CBell wrote initial manuscript draft, with contributions by LC, MKR, MJM, and RCG. All authors revised and approved the manuscript.

## Materials and Methods

### Cell Culture

HeLa cells were maintained at 37 °C in a humidified atmosphere containing 5% CO2 and cultured in Dulbecco’s Modified Eagle Medium (DMEM, Gibco) supplemented with 10% (v/v) fetal bovine serum (FBS, Gibco), 1% (v/v) penicillin–streptomycin (Sigma) and 1% (v/v) GlutaMAX (Thermo). Selective antibiotics were added according to the requirements of the expression constructs: blasticidin S (6 µg/ml), puromycin (0.5 µg/ml), hygromycin B (0.3 mg/ml). The parental HeLa cell line (‘Kyoto strain’) was obtained from S. Narumiya (Kyoto University, Japan) and validated by a Multiplex human Cell line Authentication test (MCA). All cell lines used in this study were routinely tested and confirmed to be free of mycoplasma contamination.

### Cell Line Generation

A monoclonal ΔCENP-B cell line was generated by a Cas9-induced frameshift mutation. SMC4-AID-Halo cells were seeded in a 24-well plate and transfected the following day with paired gRNAs using Lipofectamine CRISPRMAX Cas9 Transfection reagents (Thermo Scientific), according to manufacturer’s guidelines, resulting in transfection of 1.13 µg Cas9 protein with 6.73 pmol respective sgRNAs per well. Cells were incubated with transfection mix for 48 h before diluting for monoclonal expansion. ΔCENP-B clones were identified by immunofluorescence for the absence of CENP-B signal (see below). Clonal characterisation included assessment of growth rate and SMC4 auxin-dependent depletion.

Cell lines containing TetO repeats were generated by random integration. SMC4-AID-Halo cells lacking puromycin resistance were seeded in a 15 cm dish and transfected the following day with a plasmid containing 250 repeats of the TetO sequence and a puromycin resistance cassette using PEI (72 µg PEI, 18 µg DNA). After two days, stable selection was initiated with puromycin (0.5 µg/mL), and maintained for 14 days prior to manual picking of individual clones for further expansion. To visualise TetO integration, a TetR-GFP-containing plasmid was transiently transfected and clones were imaged using a confocal microscope (Zeiss LSM780). Multiple clones exhibiting GFP foci consistent with a single TetO integration pattern were further expanded and characterised.

### Condensin-degradation and microtubule depolymerisation

Prior to cell seeding, dishes were coated with 0.01% Poly-L-Lysine (Sigma) and incubated for 20 min, followed by three washes with dH2O. Cells were pre-synchronised by an overnight block in 2 mM Thymidine (Sigma) and released into S phase by washing cells three times with fresh DMEM. 6 h later, cells were treated with 10 µM RO-3306 (Sigma), 1 µM 5-Ph-IAA (Bio Academia) and 3.3 µM Nocodazole (Sigma), thus synchronising cells to the G2/M boundary, acutely degrading condensins and preventing spindle polymerisation, respectively. Cells were released into mitosis 18 h later by washing three times with fresh DMEM containing 5-Ph-IAA and Nocodazole and fixed for immunofluorescence 5 h later (see below).

#### HaloTag staining

Successful condensin-depletion was determined by staining for HaloTag in both condensin-degraded and untreated SMC4-AID-Halo cells, as well as in WT HeLa Kyoto cells without HaloTag. One hour before fixation, HaloTag TMR Ligand (Promega) was added to cells in a 1:1000 dilution in respective media. Following a 30 min incubation, cells were gently washed five times with fresh respective media and left another 30 min without ligand before fixation.

#### Mitotic and apoptotic cell staining

To confirm that the condensin-degradation and microtubule spindle depolymerisation protocol did not induce apoptosis, cells were additionally stained for cleaved-Caspase-3 using CellEvent Caspase-3/7 Green (Thermo Scientific) upon release into mitosis following manufacturer’s instructions. For control, unsynchronised cells were treated with either just CellEvent Caspase-3/7 Green, or CellEvent Caspase-3/7 Green and the apoptosis-inducing BH3-mimetics ^51^ 5 µM ABT-737 (Selleck Chemicals) and 0.5 µM S63845 (Selleck Chemicals) in parallel. All cells were fixed after 5 h and stained for phospho-H3Ser10 by immunofluorescence (see below) to additionally assess mitotic state.

### Kinetochore depletion

To deplete kinetochores, cells were seeded at very low concentration (∼6% confluency) in 0.01% Poly-L-Lysine (Sigma) coated 35 mm µ-Dishes (ibidi) in DMEM. The following day, Cas9-RNPs were generated using Lipofectamine CRISPRMAX Cas9 Transfection reagents (Thermo Scientific) according to manufacturer’s protocol, using either a custom CENP-C-targeting TrueGuide sgRNA (Thermo) or a negative control TrueGuide sgRNA (Thermo) for control conditions, resulting in transfection of 4.55 µg Cas9 protein with 28 pmol sgRNA per dish. 22 h after this, cells were washed with fresh DMEM and transfected with an siRNA pool targeting CENP-C (Dharmacon^52^) or a non-targeting siRNA (Dharmacon) using Lipofectamine RNAiMAX (Life Technologies) according to manufacturer’s protocol, adding 20 pmol siRNA to each dish. Cells were washed the next day to remove transfection reagents. 76 h after Cas9-transfection, cells were pre-synchronised in Thymidine and treated as in <u>Condensin-degradation and microtubule depolymerisation</u> (see above), with addition of 12 µM proTAME (THP medical products) alongside Nocodazole (Sigma) and 5-Ph-IAA (Bio Academia) to ensure mitotic arrest also in the absence of kinetochores and a functioning spindle-assembly checkpoint.

### Condensin-degradation and microtubule depolymerisation - RASER-FISH

To perform RASER-FISH on condensin-degraded and microtubule-depolymerised cells, cells were seeded on 8-well slides (ibidi) previously coated with 0.01% Poly-L-Lysine (Sigma) and pre-synchronised by an overnight block in 3 µg/ml Aphidicolin (Sigma) with 40 µM BrdU/BrdC (Sigma/Santa Cruz; 3:1 ratio). Cells were released into S phase by washing cells three times with fresh DMEM containing 40 µM BrdU/BrdC. After 6 h, cells were cultured overnight in DMEM containing 10 µM RO-3306 (Sigma), 3.3 µM Nocodazole (Sigma), 1 µM 5-Ph-IAA (Bio Academia) and 40 µM BrdU/C. Cells were released into mitosis 18 h later by washing three times with fresh DMEM containing 5-Ph-IAA and Nocodazole. After 5 h, cells were washed 3x with 1x PBS and fixed in 4% paraformaldehyde (PFA) in 1x PBS for 15 min at room temperature, quenched with 100 mM NH4Cl for 10 min and washed three times with 1x PBS. Cells were either directly further processed for RASER-FISH or stored at 4 °C. For conditions including TetR-GFP transfection, quenching was performed using 10 mM TRIS-HCl (pH 7.5), before proceeding with GFP-boosting (see below, <u>TetO surface targeting assay - GFP Booster</u>) prior to RASER-FISH.

### RASER-FISH

For RASER-FISH, freshly prepared buffers were used and all steps were done at room temperature unless noted. DNA was UV-sensitised by incubation with 0.5 µg/ml DAPI in 1x PBS for 15 min, PBS-washed, and irradiated at 254 nm (3.6 J/cm^2^, ∼20 min) in a crosslinker; Exonuclease III (1 U/µl in 1x NEBuffer 1) was applied for 15 min at 37 °C in a humidified chamber; cells were pre-equilibrated for 1 h at 37 °C in hybridisation buffer HB1 (10% w/v dextran sulfate, 50% v/v formamide in 2x SSC) and hybridised overnight at 42 °C in HB1 containing primary probes (∼2 nM/probe; also encoding binding sites for fluorescently labelled secondary oligos). The next day, cells were washed twice in wash buffer WB1 (50% v/v formamide in 2x SSC) at 35 °C for 5 min each, treated with RNase H (0.05 U/µl in 1x RNase H buffer) for 20 min at 37 °C, washed three times in WB2 (0.2% v/v Tween-20 in 2x SSC) for 3 min each, incubated for 3 min in HB2 (5% w/v ethylene carbonate, 0.2% v/v Tween-20, 20 nM fluorescently labelled secondary oligo in 2x SSC), washed once in WB3 (10% v/v formamide, 0.2% v/v Tween-20 in 2x SSC) for 1 min, and finally placed in imaging buffer (0.2 µg/ml DAPI in 2x SSC) for imaging.

### TetO surface targeting assay

For RASER-FISH experiments regarding TetO, cells were seeded on an 0.01% Poly-L-Lysine (Sigma) coated 8-well slide (ibidi) and transfected the following day with TetR-GFP(neg25) plasmid using XtremeGENE 9 according to manufacturer’s instructions, resulting in 70 ng plasmid per well. The following day, cells were pre-synchronised to S phase and treated as in <u>Condensin-degradation and microtubule depolymerisation - RASER-FISH</u> with minor modifications: Nocodazole was used at 0.6 µM and for relevant conditions, Doxycycline was provided from RO-3306 arrest until fixation at 1 µg/ml in all media and buffers.

### TetO surface targeting assay - GFP Booster

Following 15 min fixation with 4% paraformaldehyde and 10 min quenching with 10 mM TRIS-HCl (pH 7.5), cells were permeabilised with 0.2% Triton X-100, blocked with 2% BSA, and incubated with a 1:200 dilution of GFP booster (Chromotek) for 1 hour. Post-immunofluorescence, cells underwent a secondary fixation of 15 min with 4% paraformaldehyde and 10 min quenching with 100 mM NH4Cl. Cells were either stored at 4 °C or processed further as in <u>RASER-FISH</u>.

### Microinjection

For Benzonase injection, CENP-A-EGFP, SMC4-AID-Halo, H2B-mCherry-tagged HeLa cells were seeded at 60% confluency onto 0.01% Poly-L-Lysine (Sigma) coated 35mm µ-Dishes (ibidi). To ensure cells entered mitosis only shortly prior to injection, they were arrested at the G2/M boundary with 10 µM RO-3306 overnight, optionally with a pre-synchronisation using 2 mM Thymidine. Cells were washed 4x with and subsequently incubated in imaging media (DMEM with 10% (v/v) FBS (Gibco), 1% (v/v) penicillin–streptomycin (Sigma-Aldrich) and 1% (v/v) GlutaMAX (Gibco) but omitting phenol red and riboflavin to reduce autofluorescence) containing 0.6 µM Nocodazole to release the cells into mitosis before moving to a Zeiss LSM980 microscope with a mounted microinjection device (FemtoJet 4i (Eppendorf)) with InjectMan 4 micromanipulation device (Eppendorf)) and a custom-built incubator (EMBL), maintaining humidified 37 °C and 5% CO2.

Cells were injected using pre-pulled Femtotips injection capillaries (Eppendorf), which were loaded with 3 µl of Benzonase (> 99% purity, Merck) diluted 1:2 in 1 mg/ml Cascade Blue™-labelled 10 kDa Dextran dissolved in injection buffer (50 mM K-HEPES, pH 7.4, 5% glycerol, 1 mM Mg(OAc)2) using an Eppendorf Microcapillary Microloader (Eppendorf). Mitotic cells were injected using an injection pressure of 75 hPa and 0.12 s of injection time, with 20 hPa compensation pressure.

### Immunofluorescence (IF)

Prior to cell seeding, dishes were coated with 0.01% Poly-L-Lysine (Sigma) and incubated for 20 min, followed by three washes with dH2O. Cells were fixed at respective time-points for 5 min with 4% paraformaldehyde in 1x PBS after two washes in 1x PBS, followed by quenching with 10 mM TRIS-HCl for 3 min. Following further washes with 1x PBS, cells were permeabilised using 0.2% Triton-X in 1x PBS (PBS-T) for 5 min and then blocked with 2% BSA in PBS for 30 min before adding primary antibodies in 2% BSA. Cells were incubated with primary antibodies for either 2 h at room temperature or overnight at 4 °C, followed by 3x 5 min washes with 1x PBS before incubating with secondary antibodies and 1.62 µM Hoechst for 1 hr at room temperature. After 3x 5 min washes with 1x PBS, cells were kept in 1x PBS for fluorescent imaging.

### FISH probe design and amplification

FISH probe synthesis and hybridisation were performed as previously described (Beckwith, Ødegård-Fougner et al., 2025), incorporating RASER-FISH ^53^ and oligo pool amplification techniques^54^.

Primary FISH probes were designed using a HeLa Kyoto reference genome (hg19 with SNP correction^55^) – and with the addition of an artificial TetO repeat sequence for TetO targeting. The genomic positions of interest and the TetO repeat sequence were parsed for unique target sites using OligoMiner^56^ with the following parameters: oligo length of 36–42, ≥2 bp spacing, 42-46 °C annealing temperature (at 50% formamide concentration), 30–70% GC content, 0.90 LDA stringency, 18-k-mer filter of 5-7, and a secondary structure filter of 0.1. The filtered sequences were synthesised as unlabelled single-stranded DNA oligos containing internal barcode sequences for secondary probe binding, and flanked by primer sites for oligo pool amplification with a T7 promoter sequence. The oligo pool was synthesised by GenScript (GenTitan™ Oligo Pools) and amplified using 2x Phusion High-Fidelity PCR Master Mix (Thermo) with 0.5 μM primers, 15 ng template, and 1x EvaGreen (Biotium). Amplification was performed at 98 °C for 3 min, followed by cycling at 98 °C for 10 s, 66 °C for 10 s, and 72 °C for 15 s until near-plateau. PCR products were purified using the DNA Clean & Concentrator-25 kit (Zymo Research) and eluted in 50 μl ultrapure water. For in vitro transcription (IVT), 1.5 μg PCR product was incubated overnight at 37 °C with HiScribe T7 Quick High Yield RNA Synthesis Kit (NEB) in a 160 μl reaction containing 80 μl NTP mix, 6.25 μl T7 polymerase mix, 6.25 μl RNAsin Plus (Promega), and ultrapure water. Reverse transcription (RT) was carried out with 150 μl of unpurified ssRNA using Maxima H Minus Reverse Transcriptase (Thermo Fisher Scientific) with 21 μl 25 mM dNTPs (final 1.75 mM), 57 μl 100 μM forward primer (final 19 μM), 6 μl RNAsin Plus, 6 μl RT enzyme (1,200 U), and 60 μl 5× RT buffer at 50 °C for 1 hour. RNA was degraded by adding 150 μl 0.5 M EDTA and 150 μl 1 M NaOH, followed by incubation at 95 °C for 15 min. The resulting single-stranded DNA probes were purified using the DNA Clean & Concentrator-100 kit (Zymo Research), eluted in 200 μl ultrapure water, and quantified using a Nanodrop spectrophotometer.

The secondary probes were fluorescently labelled by click chemistry (ClickTech Oligo Link Kit, baseclick) using 3′-C3-azide oligos (Metabion) and an alkyne-modified ATTO647N fluorophore (ATTO-TEC). The labelled oligos were HPLC purified and resuspended in 1x TE buffer.

### Cell Microscopy & Image Processing

Imaging was performed on a Zeiss LSM980 microscope equipped with an incubator chamber (EMBL) using either a 40x 1.3 NA or a 63x 1.4 NA oil-immersion DIC plan-apochromat objective (Zeiss). The system was integrated with an Airyscan2 detector and operated by ZEN3.3 Blue 2020 software. Cells were imaged using confocal mode, with the exception of RASER-FISH experiments, for which a combination of Airyscan (foci detection) and confocal (chromatin detection) was used. For live cell microinjection experiments, incubation was set to maintain a humidified atmosphere and a constant 37 °C temperature with 5% CO2. For immunofluorescence experiments, chromatic aberration from the microscope was corrected by image registration using 0.2µm TetraSpeck Microspheres (Thermo Fischer Scientific, prepared in-house and spread onto glass slides) and a custom Fiji script. For RASER-FISH experiments, 3D Airyscan processing was performed for each acquired image using default parameters. Images were linearly contrast adjusted and matched across conditions for all representative microscopy images, unless stated otherwise.

### Microinjection FRAP

For Fluorescence Recovery After Photobleaching (FRAP) experiments, a single frame was photobleached in the designated region of interest using a laser intensity 200-fold greater than that used for image acquisition, while simultaneously increasing pixel dwell time from 0.86 μs to 2.84 μs. Images were captured every 5 s for the duration of the experiment, with 10 frames recorded prior to bleaching.

### Image analysis – Linescan

The distance of individual foci relative to the chromatin boundary was measured using a semi-automated analysis pipeline (see below, <u>Linescan software</u>). For unperturbed WT prometaphase cells, lines were drawn through the centre of CENP-A foci into the corresponding sister chromatid arm where these pairs could be clearly identified and were in a planar arrangement (± one z-slice). For condensin-depleted or Benzonase-injected cells, lines were drawn through the respective foci perpendicularly to the nearest chromatin surface. Where the chromatin signal intensity of inner inclusions went below 50% of the surrounding chromatin signal intensity, these boundaries were also considered as chromatin surface. Line-width was chosen to cover ∼75% of focus width and lines were drawn with sufficient length to reach a stable signal plateau both outside and inside of chromatin. Foci that were located at a chromatin surface grazing sections, such that a change of ± one z-slice significantly altered the measurement result, were excluded from the analysis. A threshold of 200 nm inside of the chromatin boundary was used to determine the fraction of foci at the chromatin surface.

### Image analysis - SMC4 degradation

Condensin degradation was measured on cells imaged using a Zeiss LSM980 in confocal mode with 1 µm z-slice interval. Maximum intensity z-projections were thresholded with Otsu dark mode in Fiji after applying a Gaussian blur filter (σ = 2) on chromatin to determine the region of interest (ROI) in which to measure mean signal intensities. The ratio of Halo/chromatin signal from cells without HaloTag was subtracted from the Halo/chromatin ratio from condensin-degraded and untreated SMC4-AID-Halo cells to account for background staining. Ratios were averaged across multiple cells in each replicate and normalised to untreated SMC4-AID-Halo cells.

### Image analysis - Microtubule depolymerisation

Successful microtubule depolymerisation of condensin-degraded and spindle depolymerised cells was assessed by staining for Tubulin by immunofluorescence (see above). Soluble Tubulin staining was used to set a threshold (Otsu) of the cell body, using the “fill holes” and “watershed” function where necessary to obtain a single ROI covering the entire cell. Tubulin mean signal intensity and standard deviation were measured within the ROI in Fiji and used to calculate the coefficient of variation per cell. Within each replicate, the mean coefficient of variation across all cells in each condition was normalised to unperturbed prometaphase cells.

### Image analysis – Mitotic and apoptotic staining

Cells were imaged using a Zeiss LSM980 in confocal mode with 1 µm z-slice interval. Mitotic and apoptotic staining was measured on maximum intensity z-projections. Measurement ROIs were determined by applying a Gaussian blur filter (σ = 2) on chromatin and thresholding in Fiji (Otsu). Mean signal intensity was measured in ROIs across all channels and four 25×25 px circles were drawn outside of cells to determine signal background. Both pH3-Ser10 and CellEvent Green signal were normalised over chromatin signal per cell and averaged by experiment round. For mitotic state, pH3-Ser10 over chromatin signal was normalised for each condition to control mitotic cells. For apoptotic state, CellEvent Green signal (over chromatin) was normalised for each condition to BH3-mimetic-treated cells.

### Image analysis – Chromatin density

The chromatin density before and after chromatin fractionation was determined by first manually selecting the central plane of the chromatin z-stack, before drawing three 6x6 px circular ROIs at random positions. All values were normalised such that the mean chromatin density of pre-injected cells was equal to 1.

### Image Analysis – Chromatin mobility by FRAP

Chromatin mobility as assessed by FRAP, with measurements performed according to ^57^. In short, rigid-body registration was performed using TurboReg. Chromatin fluorescence signal was measured across three ROIs (FRAP, total chromatin and background) per time-lapse movie using the Time Series Analyzer v.3 (J. Balaji): The FRAP ROI was manually drawn as a 10x10 px circle in the bleach region, while the background ROI was manually drawn as a 25×25 px circle away from chromatin. To determine the total chromatin ROI, a gaussian blur filter (σ = 2) was applied to chromatin in the first recorded frame, followed by thresholding using the Intermodes dark method in Fiji.

### Image Analysis – Kinetochore depletion

To confirm depletion of various kinetochore proteins (CENP-C, NDC80, KNL1), 6x6 px circular ROIs were manually drawn on CENP-A foci used in the localisation quantification (linescan tool) and the mean intensity of the respective kinetochore protein measured. Fluorescence intensity was normalised across the individual z-stack and cell by manually determining the quartile z-slices of the chromatin mass using the Hoechst chromatin stain, before drawing four 15×15 px circles on chromatin away from CENP-A on these quartile slices and extracting mean intensity for chromatin and respective kinetochore protein signal. The mean signal across circles across quartile slices for kinetochore signal was subtracted as background from centromeric ROIs. ROIs were each normalised over the mean chromatin signal per cell. ROIs were further normalised between conditions across each biological replicate.

### ΔCENP-B cell line validation

Fields of views for ΔCENP-B and control cells were imaged after immunofluorescence staining for CENP-A and CENP-B. The absence of CENP-B signal at centromeres in ΔCENP-B cells was quantified by thresholding CENP-A with Otsu dark mode in Fiji after applying a Gaussian blur (σ = 2) and extracting the mean fluorescent signal of CENP-A and CENP-B within this selection. Background signal for both protein channels was determined by drawing a 15×15 px circle in five cells on chromatin (stained by Hoechst) away from CENP-A. Following background subtraction, the ratio of CENP-B over CENP-A signal was calculated and normalised to control cells.

### Linescan software – Overview

We developed a Python-based linescan analysis tool that quantifies spatial relationships between fluorescence signals along user-defined line segments. For each region of interest (ROI), the software extracts two one-dimensional intensity profiles: a reference profile used to define the spatial zero and a target profile used to localise a peak relative to that zero. The pipeline registers profiles by estimating the first half-maximum crossing of the reference profile via a logistic (sigmoid) fit with robust least squares (soft_l1) and a residual-based fallback to polynomial smoothing; the target peak is estimated by a baseline-plus-Gaussian fit with a residual-based fallback to a polynomial method. In typical applications, the sigmoid alignment is applied to the chromatin channel to detect the chromatin surface, whereas the Gaussian peak model is used for compact, spot-like signals such as immunofluorescence or FISH foci. The pipeline operates deterministically on each ROI independently.

### Linescan software - Image data and ROI definition

Images are multi-channel stacks (e.g., multi-plane TIFFs). ROIs are provided as ImageJ/Fiji line ROIs in.roi or.zip format. For each ROI, we retrieve the line endpoints and slice metadata and use a profile width (default 5 pixels) to average intensities orthogonally to the line segment, yielding a single 1D signal per channel. For each ROI, the tool extracts 1D profiles for the reference and target signals, with the x-axis in pixel indices subsequently converted to physical units using a user-supplied pixel size scaling factor (e.g., micrometers per pixel).

### Linescan software - Alignment offset by half-maximum of a Sigmoid fit

To register profiles across ROIs, we estimate an alignment offset on the reference signal as the position of the half-maximum on the rising edge. By default, we fit a four-parameter logistic (sigmoid) model to the raw reference profile using robust nonlinear least squares (soft_l1; covariance disabled with calc_covar=False):

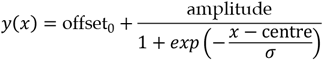

The fitted centre parameter gives the exact half-maximum crossing and is taken as the alignment offset. For visualisation, the fitted curve is evaluated on a dense grid. Residual quality is quantified by normalised RMSE (relative to the data’s dynamic range) and R^2^; if quality is poor (defaults NRMSE ≤ 0.12 and R^2^ ≥ 0.75), or if the fit fails to converge, we fall back to polynomial smoothing (degree ≤ 10, limited by profile length), evaluate on a dense grid, compute the half-maximum level, and locate the first rising crossing by linear interpolation.

### Linescan software - Peak localisation by Gaussian modelling

Peak positions in the target profile are estimated by fitting a Gaussian peak on a constant baseline to the raw profile:

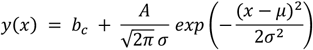

where A is the Gaussian area parameter used by lmfit. Initialisation is robust: the baseline is set to the 10th percentile; the detrended profile is smoothed with a Savitzky–Golay filter (odd window ≈ 40% of profile length, polyorder ≤ 3) to stabilise peak detection; up to three candidate peak centres are identified by prominence (threshold 10% of the dynamic range); widths are initialised from half-maximum widths and converted via FWHM = 2.355σ. Parameters are constrained with reasonable bounds and amplitude capping. Fitting uses robust least squares (soft_l1). The Gaussian centre μ from the best fit *is* taken as the peak position. Residual quality is evaluated with NRMSE and R^2^ (defaults NRMSE ≤ 0.12, R^2^ ≥ 0.70); if quality is poor or the fit fails, we fall back to a degree-capped polynomial evaluated densely and report the tallest peak using a height-based peak finder (threshold ∼60% of the smoothed maximum).

### Linescan software - Distance computation, normalisation, and visualization

For each ROI, the aligned distance between the peak and the half-maximum is computed as (μ−offset) and converted to physical units using the calibrated pixel size. Raw profiles are not normalised prior to fitting; min–max normalisation is applied only for display overlays. Where indicated, we plot raw and fitted curves for both signals on a common x-axis shifted by the alignment offset.

### Linescan software - Implementation details

The tool is implemented in Python using NumPy and SciPy (signal processing and optimisation), scikit-image (I/O and profile extraction), lmfit (nonlinear model fitting), Matplotlib and Seaborn (visualisation), and the read-roi library for parsing ImageJ/Fiji ROI files. The complete linescan workflow includes ROI parsing, profile extraction, alignment, model fitting, and plotting and summary visualisations.

### Linescan software - Reproducibility and availability

All figures in this study were generated using this software together with companion Jupyter notebooks available at GitHub: https://github.com/gerlichlab/bell_et_al. The analysis tool described here is openly available at GitHub: https://github.com/gerlichlab/linescan. The software produces a per-ROI results table and reproducible figures from the same inputs and parameters, enabling end-to-end reproducibility from raw images and ROIs to the reported measurements and plots.

### Generation of minimal *in silico* chromatin and protein models

For our in-silico study, we employ our multiscale coarse-grained chromatin model that integrates information at three complementary resolution levels: (i) atomistic data of nucleosomes, histone proteins, and DNA, (ii) a DNA and amino acid chemically-specific sequence-dependent model, and (iii) a minimal chromatin model, deployed here, that represents each nucleosomes with just ∼30 particles^44,45^. Nucleosome–nucleosome and nucleosome–DNA interactions in this minimal model are parametrised via PMFs calculated with our higher resolution, chemically-specific model.

For each nucleosome, a central bead represents the histone octamer, and the rest are used to model the surrounding DNA. Each bead representing the DNA signifies five base pairs and is classified as either nucleosomal DNA, which is permanently and rigidly bound to the nucleosome core, or linker DNA, which is free to move and connected via bonded interactions. These interactions, which crucially capture the mechanical rigidity of the DNA, are described with a coarser version of the Rigid Base-Pair (RBP) model that also underpins our chemical-specific resolution. For a complete description of this parametrisation, please refer to ^44^.

Each bead, both for the nucleosome core and DNA, is a spherical rigid body described by a position vector, 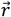, and a unit quaternion, 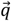, which represents its relative orientation to surrounding beads. The DNA-bonded interaction uses an RBP-like potential^44,58,59^, which has a functional form depending on the six helical parameters of DNA:

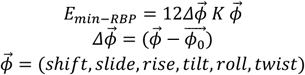

Here 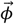 is a 6-dimensional vector representing all helical parameters, or independent degrees of freedom, between two DNA basepairs, or in this case, two beads. These are three angles (roll, tilt and twist) and three distances (shift, slide and rise). 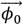 corresponds to the equilibrium value of these helical parameters, and K is the 6x6 stiffness matrix. Both 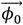 and *K* are parametrised from higher resolution MD simulations, as described in ^44^. The equilibrium rise between base pairs is 16.45 A°, the equilibrium twist is 172 degrees, and the equilibrium values of the other helical parameters are zero. We set *K* to be a fully diagonal stiffness matrix with diagonal values:

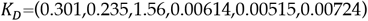

Pairwise interactions are mapped onto shifted and truncated Lennard-Jones (LJ) potentials, that closely match the pair-force matching profiles in our higher resolution model. DNA-DNA interactions are purely repulsive, similar to the electrostatic repulsion of negatively charged DNA. In contrast, DNA-Core interactions are attractive, resembling the electrostatic attraction between DNA and histone core proteins. The interaction strength *ϵ* is empirically related to the monovalent salt concentration, following an established mapping (found in ^44^). The total energy of the system is then

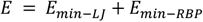

where *E*_*min−LJ*_ is the sum of all LJ terms, and *E*_*min−RBP*_ is the sum of all bonded RBP-like terms. Simulations are performed using Langevin dynamics in a custom modified version of LAMMPS ^60^, with solvent effects included via a mean-field approximation to electrostatic screening. This approach presumes that salt concentration primarily influences the screening of the mean electrostatic potential, simplifying the model while still capturing the essential LLPS behaviour. It overlooks ion-specific effects like correlations, polarisation, release upon binding, and diverse binding patterns. These effects are likely minor under monovalent, physiological conditions but could become more important with polyvalent ions or elevated salt levels.

In this study, we expand the minimal chromatin model to include coarse-grained representations of CENP-B and TetR–GFP proteins. These proteins are modelled at the same resolution as chromatin: the globular domain of the CENP-B protein is represented as a sise-consistent, large spherical bead. Its long disordered domain, which in atomistic reality carries a net negative charge, is modelled as a flexible polymer chain whose beads interact with one another and with DNA through repulsive interactions. These interaction strengths are assigned using the same empirical mapping between overall composition and bead– bead potentials as for DNA, ensuring that the domain’s net tendency for repulsion is captured in a coarse-grained, mean-field fashion. In this representation, the disordered domain exhibits scaled repulsion with itself and DNA, and a scaled attraction to the histone core. Similarly, for the TetR–GFP(neg25) construct, the TetR dimer—an ordered domain with high affinity for the Tet-O sequence placed in the ligated DNA—is represented as a single bead bound to the ligated DNA. The attached GFP(neg25) motif, which in atomistic reality carries a net negative charge, is represented as a second large bead whose bead–bead interaction parameters are assigned to produce self-repulsive behaviour and repulsion from DNA, following once again the same salt-dependent empirical mapping used for the CENP-B disordered domain.

### Creating initial *in silico* structures

To generate physically accurate chromatin array structures, we start with the chemically specific model to construct an initial configuration that has the desired linker DNA length at a 1 bead-per-base-pair resolution. Next, we convert this structure into the minimal model by grouping every five DNA base pairs into one bead; any leftover base pairs at the end of the array are then discarded.

For simulations involving CENP-B or TetR–GFP, the proteins are initially placed directly onto their specific DNA binding sites, found in the ligated DNA at one end of the array. Given their strong binding and long residence times, we assume these proteins stay bound throughout the simulation period, as the simulated timescales are short relative to experimental dissociation times and the goal here is to probe the system’s structural organisation under stably bound conditions. Occupancy is set at two-thirds of the available binding motifs, that is, as experimental ligated DNA constructs have three motifs per site, we assume occupancy of two of them.

### Single fibre MD simulation details

All single fibre MD simulations in this study were performed in LAMMPS^61^ using Langevin dynamics and temperature replica exchange molecular dynamics (T-REMD) in the NVT ensemble, with replicas spanning 300– 600 K. The number of replicas was optimised to the system size to maintain exchange probabilities near 0.3. Each run lasted at least 200 million timesteps, with exchange attempts every 100 timesteps. Snapshots of coordinates were recorded every 100,000 timesteps. The first half of each trajectory was discarded, and data analysis focused only on the 300 K replicas.

### Direct Coexistence MD simulations

To probe the surface targeting properties of each construct and the overall organisation of the mixed system condensates, we simulated 120 fibre chromatin systems at a fixed monovalent salt concentration. The modelling protocol was adapted to handle mixtures of fibre types, with 10% of the fibres (12 out of 120) designated as modified arrays and the remainder as canonical arrays. Initial configurations for individual fibres were generated from T-REMD simulations at room temperature and the target salt concentration. Here, the target salt was set to 0.075 M. Under these conditions, Direct Coexistence simulations^62–64^ (described below) of chromatin solutions using our minimal coarse-grained model produce an equilibrium chromatin-rich condensed phase in coexistence with a chromatin-poor diluted phase. We have shown that the stability and material properties of these simulated chromatin condensates are in agreement with those measured in vitro^45^. In the Direct Coexistence method, condensed and dilute chromatin phases are placed. In the same elongated rectangular 3D box (dimensions 1,200 Å × 1,200 Å × 6,000 Å) separated by an interphase. MD simulations are then performed in the NVT ensemble, and after equilibrium is reached, uncorrelated frames are used to perform statistical analyses. Uncorrelated frames were obstained every 50,000 timesteps (1 timestep = 100 fs; correlation time ≈ 4,000 timesteps). All these simulations were performed in LAMMPS^61^ using Langevin dynamics.

These simulations focused on analysing the internal molecular organisation of chromatin arrays of different types within the same condensate. We observed that layered condensates arise from chromatin solutions combining canonical chromatin arrays with chromatin arrays that present electrostatic polarity (e.g. arrays with either CENP-B, 300 or 500-bp DNA, or TetR-GFP(neg25) ligated to one end). To understand the internal structure of condensates, we computed one-dimensional density profiles along the long axis of the simulation box. For each uncorrelated simulation frame, the box was segmented into bins along this axis, and the number density of each fibre type (canonical or modified) was determined per bin. Average density profiles were generated from the equilibrated portion of each simulation after skipping 9 out of every 10 saved frames to reduce data size while maintaining statistical independence. This downsampling yielded datasets of ∼500 frames per simulation, compared to the original ∼5,000 uncorrelated frames. The remaining frames were divided into 20 blocks, each containing ∼25 independent frames, and a density profile was computed for each block. The resulting block-averaged profiles were then combined to obtain the mean and standard deviation used to characterise the layered organisation and its variability. For clarity, individual arrays in the dilute phase are removed from slabs shown in figures.

### *In silico* surface enrichment analysis

To compute the preference of modified fibres to localise to either the interface or the bulk of the condensate, we computed symmetrised density profiles along the simulation box’s long axis. The box was segmented into bins, and for each, we calculated the total fibre density and the density of modified fibres. Profiles were block averaged as described above, and symmetrised by taking the absolute coordinate relative to the condensate centre. The interface position was identified as the point of maximum gradient in the total density profile. An interface region was then defined by extending five bins on each side of this point (11 bins total, corresponding to ∼44 nm along the box axis), a choice that balances capturing the interfacial zone while minimising inclusion of bulk regions. The number of bins was determined empirically by visual inspection of density profiles to ensure that the defined region fully encompassed the interface without overlapping excessively with the bulk. The interface enrichment in composition was calculated as the ratio of the fraction of modified fibres in the interface region to that in the bulk:

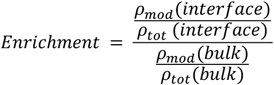

where *ρ*_*mod*_(*interface*) is the average density of modified fibres in the interface region, *ρ*_*tot*_(*interface*) is the average density of total fibres in the interface region, *ρ*_*mod*_(*bulk*) is the average density of modified fibres in the bulk region and *ρ*_*tot*_(*bulk*) is the average density of total fibres in the bulk region. A compositional enrichment of 1 indicates that the fraction of modified fibres at the interface is the same as in the bulk, meaning no preferential localisation. Values greater than 1 indicate enrichment at the interface (e.g., 10 means tenfold higher proportion at the interface), while values less than 1 indicate depletion from the interface. This process was repeated for each data block to generate blockwise enrichment values, which were then summarised as distributions for statistical analysis and represented here in boxplots showing the 5-95 percentile, not showing outliers.

### PMF calculations

We calculated PMFs from umbrella sampling MD simulations with the COLVARS module^65^. To enhance sampling efficiency and capture the broad spectrum of chromatin conformations at a fixed collective variable (CV), we implemented temperature replica exchange molecular dynamics (T-REMD) within each umbrella window. Each window was run independently without exchanges between different CVS; instead, temperature swaps occurred among 16 replicas ranging from 300 to 600 K, following the Metropolis acceptance criterion, with the umbrella bias incorporated into the total potential energy.

For fibre–fibre PMFs, the CV was the distance between the centres of mass of two chromatin fibres, biased by a harmonic potential with a force constant of 0.002 kcal mol^−1^ Å^−2^. Sixteen equally spaced windows spanned from 0 to 600 Å.

For fibre–bulk PMFs, we chose a single chromatin fibre within the condensed phase and defined the CV as the distance between its centre of mass and that of the remaining fibres (the “bulk”). PMFs were reconstructed using the Weighted Histogram Analysis Method (WHAM, http://membrane.urmc.rochester.edu/wordpress/?page_id=126) from the combined data of the replicas.

### Structure analysis

The radius of gyration *R_g_* was used to evaluate the structural compactness of each simulated fibre. For each saved simulation frame of an equilibrated single-molecule replica exchange, *R_g_* was determined as the root mean square distance of all beads from the centre of mass of the fibre:

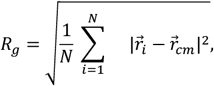

where N is the total number of beads, 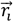 the position vector of bead *i* and 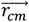 the centre of mass of the array, computed as

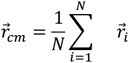

This measure captures the overall spatial distribution of the fibre and provides a single value describing its compactness. To minimise the influence of early nonequilibrium conformations, only the second half of each trajectory was used in the analysis. For every system presented here, *R_g_* values were extracted directly from LAMMPS dump files, stored in per-frame CSV files, and then plotted. The plots correspond to the *R_g_* distribution of ∼5000 different frames.

### Calculation of acidic segments in the human proteome

The complete human proteome of all reviewed (Swiss-Prot) canonical proteins (20,405 entries) was obtained from UniProt. Each protein sequence was computed with variable-sized sliding windows to identify a segment with the lowest net charge. Two scoring systems were applied:

1. Aspartic acid (D) and glutamic acid (E) were assigned –1, arginine (R) and lysine (K) were assigned +5, and all other residues were assigned a gap penalty of 0.1.
2. Aspartic acid (D) and glutamic acid (E) were assigned –1, arginine (R) and lysine (K) were assigned +5, and all other residues were assigned a gap penalty of 0.5.

For both scoring systems, CENP-B ranked second in the human proteome for containing the longest negatively charged segment.

### Preparation of 12×601 DNA array for *in vitro* chromatin assembly

The bacterial expression vector containing 12 repeats of Widom’s 601 sequence^66^ and a 37 bp extension (described below) was cloned into the pWM vector as previously described^45^. Following large-scale purification using Plasmid Giga Kit (Qiagen), the 12×601 array was liberated from the pWM vector by EcoRV (NEB), and the resulting blunt ends were dephosphorylated by Quick CIP (NEB)^27,28,45^. The DNA mixture was purified by phenol-chloroform extraction and precipitation. The 37 bp extension on the 12×601 array contains a recognition site for BbsI, a type IIS restriction enzyme that cleaves outside of its recognition sequence. By design, digestion of the 12×601 array bearing the 37 bp extension by BbsI (NEB) generated an overhang of GGGG. The mixture of BbsI-digested 12×601 array and cleaved pWM vector was purified by another round of phenol-chloroform extraction and precipitation.

### Preparation of ligation-competent DNA fragments containing CENP-B-boxes or TetO

Synthetic single-stranded oligonucleotides were purchased from IDT. One strand carried a 5′ phosphate and a CCCC extension, while the complementary strand carried a 5′ Cy5 label. Oligonucleotides were diluted to 100 μM in Annealing Buffer (10 mM Tris-HCl pH 7.5, 1 mM EDTA pH 7.5, 50 mM NaCl). Complementary strands were annealed by heating to 99 °C followed by slow cooling to room temperature. The resulting double-stranded DNA contained a 5′ Cy5 label at one end, a 5′ phosphorylated CCCC overhang at the other end, and either four CENP-B-boxes (CTTCGTTGGAAACGGGA) or three TetO sequences (TCCCTATCAGTGATAGAGA).

### Preparation of ligation-competent 100 bp, 300 bp and 500 bp DNA

DNA sequences were amplified from an unrelated 15 kb plasmid pcDNA5-BRD4-p300 (D1399Y)^67^ with the reverse primer (5’-Alexa Fluor 647-TATGTCTAGTGTACTCTGTGAGAGGTTTGAATTC) together with one of the following forward primers: GGGGTAGTCTTCGGACCTGGGACTCAGCAC, GGGGTAGTCTTCTCCAGTCCTTCCCCAAGGA, or GGGGTAGTCTTCAGGCCTATCAGCAGCGAC. The PCR reactions generated fragments of 79 bp, 279 bp, and 479 bp, respectively. The fragments were purified using the QIAquick PCR Purification Kit (Qiagen), digested with BbsI (NEB), and purified twice with the QIAquick PCR Purification Kit.

### Preparation of histone proteins and assembly of 12×601 nucleosome arrays

Expression, purification, and assembly of histone octamers and dimers containing *H. sapiens* histone H2A, H3 C111A, H4, and H2B T116C (unlabelled, Alexa Fluor 488–labelled, or Alexa Fluor 594–labelled) were performed as previously described^45^. Nucleosome arrays were assembled from 12×601 DNA and histone proteins following the same method ^45^.

### Preparation of 12×601 nucleosome arrays with extensions containing CENP-B-boxes, TetO or nucleosome-free DNA fragments

12×601 nucleosome arrays containing 5’ phosphorylated GGGG overhangs were ligated to DNA fragments containing 5’ phosphorylated CCCC overhangs and 4x CENP-B-boxes, 3x TetO sequences, 79 bp, 279 bp, or 279 bp DNA. Ligation reactions were performed at a final concentration of 83 nM nucleosome array-DNA fragment pairs in 2 mM MgCl2, 50 mM Tris-HCl pH 7.5, 1 mM ATP, 10 mM DTT, and 300 units/μl T7 DNA ligase (NEB). After incubation for 1 hour at 24 °C, coupled arrays were purified by a 15%-40% sucrose gradient, dialyzed, and concentrated.

### Preparation of vectors for bacterial expression of recombinant proteins

Codon-optimized sequences for *Escherichia coli* expression encoding human CENP-B (1-599), TetR and GFP(neg25)^48^ were purchased from Twist Bioscience.

The CENP-B sequence was cloned into the pMTTH vector (Gibson et al., 2019) by NdeI and NotI digestion to generate pMTTH-CENP-B. The vector is designed to express a fusion protein under the control of the isopropyl β-D-1-thiogalactopyranoside (IPTG)-inducible tac promoter. The resulting fusion protein contains CENP-B flanked at the N-terminus by a maltose-binding protein tag (MBP) and a TEV protease recognition site (MT), and at the C-terminus by a TEV protease recognition site followed by a 6xHis tag (TH). To generate deletion constructs, the acidic region (residues 404-538) and the dimerization domain (residues 539-598) were removed by PCR amplification of the flanking plasmid followed by ligation, yielding pMTTH-CENP-B (Δacidic) and pMTTH-CENP-B (Δdimer), respectively.

To generate pM33H-TetR-GFP(neg25), the TEV protease recognition sites in pMTTH were replaced with the HRV-3C protease recognition site, and TetR and GFP(neg25) were inserted by Gibson assembly.

### Recombinant protein expression and purification

All proteins of interest were expressed and purified following a similar method. *Escherichia coli* strain BL21(DE3) cells harboring protein expression vectors (e.g., pMTTH-CENP-B, pM33H-TetR-GFP(neg25)) were grown in Luria broth supplemented with 100 µg/mL ampicillin at 37 °C until reaching an OD600 of 0.8. Protein expression was induced with 1 mM IPTG, followed by incubation for 18 hours at 18 °C.

Cells were harvested by centrifugation and resuspended in Lysis Buffer (10 mM imidazole pH 7, 50 mM HEPES pH 7, 1 M NaCl, 10% glycerol, 5 mM β-mercaptoethanol, 1 mM benzamidine, 100 µM leupeptin, 100 µM antipain, 1 µM pepstatin, 1 mM PMSF). All subsequent steps were performed at ∼4 °C. Cells were lysed by passing through Avestin Emulsiflex-C5 homogenizer once at ∼5,000 PSI and three times at ∼10,000 PSI. The lysates were clarified by centrifugation in a Sorvall RC 6 centrifuge with a SS-34 rotor at 19,500 rpm. The clarified lysates were applied to a Ni-NTA agarose resin (Qiagen) pre-equilibrated with Lysis Buffer in a gravity column and rotated for 15 minutes. The resin was washed with 20 column volumes (CV) of Nickel Wash Buffer (25 mM Imidazole, pH 7, 50 mM HEPES, pH 7, 1 M NaCl, 10% glycerol, 5 mM β-mercaptoethanol, 1 mM benzamidine, 100 µM leupeptin, 100 µM antipain, 1 µM pepstatin, 1 mM PMSF). Bound proteins were eluted with 10 CV of Nickel Elution Buffer (300 mM Imidazole, pH 7, 50 mM HEPES, pH 7, 1 M NaCl, 10% glycerol, 5 mM β-mercaptoethanol, 1 mM benzamidine, 100 µM leupeptin, 100 µM antipain, 1 µM pepstatin, 1 mM PMSF). Protein-containing fractions, detected by the Bradford assay, were applied to amylose resin (NEB) pre-equilibrated in Nickel Elution Buffer and rotated for 45 minutes. For MTTH CENP-B, MTTH CENP-B (Δdimer), and M33H TetR-GFP (neg25), the resin was washed with 20 CV of Amylose Wash Buffer (20 mM Imidazole, pH 7, 150 mM NaCl, 10% glycerol, 1 mM DTT, 1 mM Benzamidine) and eluted by 10 CV of Amylose Elution Buffer (3% maltose, 20 mM Imidazole, pH 7, 150 mM NaCl, 10% glycerol, 1 mM DTT, 1 mM Benzamidine). For MTTH CENP-B (Δacidic), the resin was washed with 20 CV of Amylose Wash Buffer (20 mM Imidazole, pH 7, 1000 mM NaCl, 10% glycerol, 1 mM DTT, 1 mM Benzamidine) and eluted by 10 CV of Amylose Elution Buffer (3% maltose, 20 mM Imidazole, pH 7, 1000 mM NaCl, 10% glycerol, 1 mM DTT, 1 mM Benzamidine).

Eluted fractions containing the protein of interest, identified by SDS–PAGE, were pooled. For MTTH CENP-B, MTTH CENP-B (Δdimer), and MTTH CENP-B (Δacidic), the MBP and 6xHis affinity tags were separated from CENP-B by incubation with TEV protease for 20 h at 4 °C. For M33H TetR-GFP (neg25), MBP and 6xHis affinity tags were separated from TetR by incubation with HRV-3C protease for 20 hours at 4°C. The MTTH CENP-B mixture was purified by a Source 15Q anion exchange column on a Fast Protein Liquid Chromatography system and eluted over a linear salt gradient from 20 mM HEPES, pH 7, 5% glycerol, 0.5 mM TCEP to 1 M NaCl, 20 mM HEPES, pH 7, 5% glycerol, 0.5 mM TCEP. Fractions containing CENP-B were concentrated and further purified by size-exclusion chromatography on a HiLoad Superdex 200 column, eluted in 20 mM HEPES pH 7, 150 mM NaCl, 5% glycerol, 0.5 mM TCEP. The MTTH CENP-B (Δdimer) mixture was purified by a Source 15Q anion exchange column on a Fast Protein Liquid Chromatography system and eluted over a linear salt gradient from 20 mM HEPES, pH 7, 5% glycerol, 0.5 mM TCEP to 1 M NaCl, 20 mM HEPES, pH 7, 5% glycerol, 0.5 mM TCEP. The MTTH CENP-B (Δacidic) mixture was purified by size-exclusion chromatography on a HiLoad Superdex 200 column, eluted in 20 mM HEPES pH 7, 1000 mM NaCl, 5% glycerol, 0.5 mM TCEP. The M33H TetR-GFP (neg25) mixture was purified by a Source 15Q anion exchange column on a Fast Protein Liquid Chromatography system and eluted over a linear salt gradient from 20 mM HEPES, pH 7, 5% glycerol, 0.5 mM TCEP to 1 M NaCl, 20 mM HEPES, pH 7, 5% glycerol, 0.5 mM TCEP.

All purified proteins were concentrated, quantified by NanoDrop, aliquoted, flash frozen in liquid nitrogen, and stored at -80 °C.

### Chromatin array phase separation experiments

Phase separation assays were performed similarly as previously described^27,28,45^ with modifications as indicated below.

#### Mixing of H3 and CENP-A arrays

83 nM 12×601 H3 nucleosome array and 83 nM 12×601 CENP-A nucleosome array were first mixed in Chromatin Dilution Buffer (25 mM Tris-OAc, pH 7.5, 0.1 mM EDTA, 5 mM DTT, 0.1 mg/mL BSA, 5% glycerol). Phase separation was then induced by addition of an equal volume of 2x Phase Separation Buffer (360 mM KOAc, pH 7.5, 6 mM Mg(OAc)2, pH 7.5, 25 mM Tris-OAc, pH 7.5, 0.1 mM EDTA, 5 mM DTT, 0.1 mg/mL BSA, 5% glycerol, 2 mg/mL glucose oxidase, 350 ng/mL catalase, 4 mM glucose, 2 µM Hoechst 33342). The phase-separated mixtures were transferred to a PEGylated microscopy plate for imaging.

#### Experiments pertaining to CENP-B

83 nM 12×601 H3 nucleosome array, 8.3 nM 4xCENP-B-Box H3 12×601 nucleosome array, and 1 μM CENP-B variant were first mixed in Chromatin Dilution Buffer (25 mM Tris-OAc, pH 7.5, 0.1 mM EDTA, 5 mM DTT, 0.1 mg/mL BSA, 5% glycerol). Phase separation was then induced by addition of an equal volume of 2x Phase Separation Buffer (200 mM KOAc, pH 7.5, 4 mM Mg(OAc)2, pH 7.5, 25 mM Tris-OAc, pH 7.5, 0.1 mM EDTA, 5 mM DTT, 0.1 mg/mL BSA, 5% glycerol, 2 mg/mL glucose oxidase, 350 ng/mL catalase, 4 mM glucose, 2 µM Hoechst 33342). The phase-separated mixtures were transferred to a PEGylated microscopy plate for imaging. Excess competitor experiment was performed by adding 500 nM free 4xCENP-B-box DNA fragment to an equilibrated, phase-separated mixture of 42 nM 12×601 H3 nucleosome arrays, 4.2 nM 4xCENP-B-Box 12×601 nucleosome arrays, and 1 μM CENP-B.

#### Experiments pertaining to nucleosome arrays fused to nucleosome-free DNA fragments

83 nM 12×601 H3 nucleosome arrays and 4.2 nM 12×601 H3 nucleosome arrays fused to nucleosome-free DNA fragments of various lengths were first mixed in Chromatin Dilution Buffer (25 mM Tris-OAc, pH 7.5, 0.1 mM EDTA, 5 mM DTT, 0.1 mg/mL BSA, 5% glycerol). Phase separation was then induced by addition of an equal volume of 2x Phase Separation Buffer (200 mM KOAc, pH 7.5, 4 mM Mg(OAc)2, pH 7.5, 25 mM Tris-OAc, pH 7.5, 0.1 mM EDTA, 5 mM DTT, 0.1 mg/mL BSA, 5% glycerol, 2 mg/mL glucose oxidase, 350 ng/mL catalase, 4 mM glucose, 2 µM Hoechst 33342). The phase-separated mixtures were transferred to a PEGylated microscopy plate for imaging.

#### Experiments pertaining to TetR-GFP(neg25)

83 nM 12×601 H3 nucleosome array, 8.3 nM 3xTetO 12×601 H3 nucleosome array, and 200 nM TetR-GFP(neg25) were first mixed in Chromatin Dilution Buffer (25 mM Tris-OAc, pH 7.5, 0.1 mM EDTA, 5 mM DTT, 0.1 mg/mL BSA, 5% glycerol). Phase separation was then induced by addition of an equal volume of 2x Phase Separation Buffer (300 mM KOAc, pH 7.5, 6 mM Mg(OAc)2, pH 7.5, 25 mM Tris-OAc, pH 7.5, 0.1 mM EDTA, 5 mM DTT, 0.1 mg/mL BSA, 5% glycerol, 2 mg/mL glucose oxidase, 350 ng/mL catalase, 4 mM glucose, 2 µM Hoechst 33342). The phase-separated mixtures were transferred to a PEGylated microscopy plate for imaging. 4 μM doxycycline hyclate was added to an equilibrated, phase-separated mixture.

All *in vitro* images shown in this study were linearly contrast adjusted to visualise respective signal over background. Figures displaying the 10:1 chromatin ratios were labelled with rounded up values, 90% and 10%.

### *In vitro* edge enrichment analysis

To determine the ratio of material in the edge region of condensates to the interior, each image was first segmented by a simple thresholding method. Briefly, each image was smoothed by a Gaussian filter, a threshold was determined by Otsu’s method, and connected pixels above this threshold were grouped together as a single segment. Next, a watershed algorithm (https://scikit-image.org/docs/0.25.x/api/skimage.segmentation.html#skimage.segmentation.watershed) was applied to dissociate segments that contained more than one condensate. Finally, segments were filtered based on size, eccentricity, and proximity to the edge of the image. In all cases, both the signal from a DNA dye (DAPI/Hoechst) and from the ligated chromatin arrays was segmented. Each segment in the DNA dye channel was the associated to a segment in the ligated array channel by proximity and a requirement of signal overlap. To account for small amounts of drift during image acquisition, paired segments from each channel were then aligned by their respective centres of mass. Following segmentation, the average fluorescent signal from both the DNA dye and ligated array of each condensate was determined as a function of pixel radius from the centre of mass, using background subtracted images to account for non-uniformity in the illumination field. Each of these radial scans began 20 pixels outside of the segment and proceeded until there were no more than 30 pixels (∼1.2 µm) remaining in the segment. The boundary of the edge was then defined as the minimum of the second derivative of the scan from the DNA channel. To reduce signal overlap, the interior was defined as beginning a further 4 pixels (∼200 nm) towards the centre of the condensate. The integrated intensity from the ligated array channel in the edge and interior regions were normalised by the respective integrated intensity from the DNA channel. The ratio of these two values was reported as surface enrichment and represented here in boxplots showing the 5-95 percentile, not showing outliers.

### Cryo Electron Tomography (cryo ET)

To prepare samples for freezing, chromatin ligated with 300 bp DNA and unligated chromatin were mixed with 2x phase separation buffer (40 mM Tris-OAc pH 7.5, 300 mM KOAc, 0.2 mM EGTA, 10%glycerol) to final concentrations of 0.166 μM and 2 μM, respectively. 4 μl of each sample was applied to glow-discharged EM grids and subjected to high-pressure freezing, as described in ^68^. The frozen grids were then imaged using a Leica cryo-fluorescence microscope to guide lamella preparation with an Aquilos FIB (Focussed Ion Beam)-SEM (Scanning Electron Microscope) (Thermo Fisher Scientific).

Data acquisition was performed on a Titan Krios G3 cryo electron microscope (Thermo Fisher) equipped with a cold-field emission gun, a Selectris X energy filter, and a Falcon 4i camera. Tilt series were collected from −48° to +60° in 2° increments, at a pixel size of 0.1516 nm, with defocus values ranging from −3 μm to −4.5 μm, and a total electron dose of 178 e−/Å^2^.

Motion correction of raw cryo-EM images was carried out using the preprocessing program Warp ^69^, followed by tilt-series alignment in AreTomo (Alignment and Reconstruction for Electron Tomography ^70^). Tomograms were reconstructed and denoised using Warp^69^ and the deep learning-based software package IsoNet ^71^. Nucleosomes were identified with context-aware template matching (CATM) as described in ^68^. DNA was manually traced in the multi-dimensional image viewer for Python, Napari, to generate 3D segmentations. Figures were created using the molecular and volume visualisation software ChimeraX ^72^.

---

**Fig. S1.**
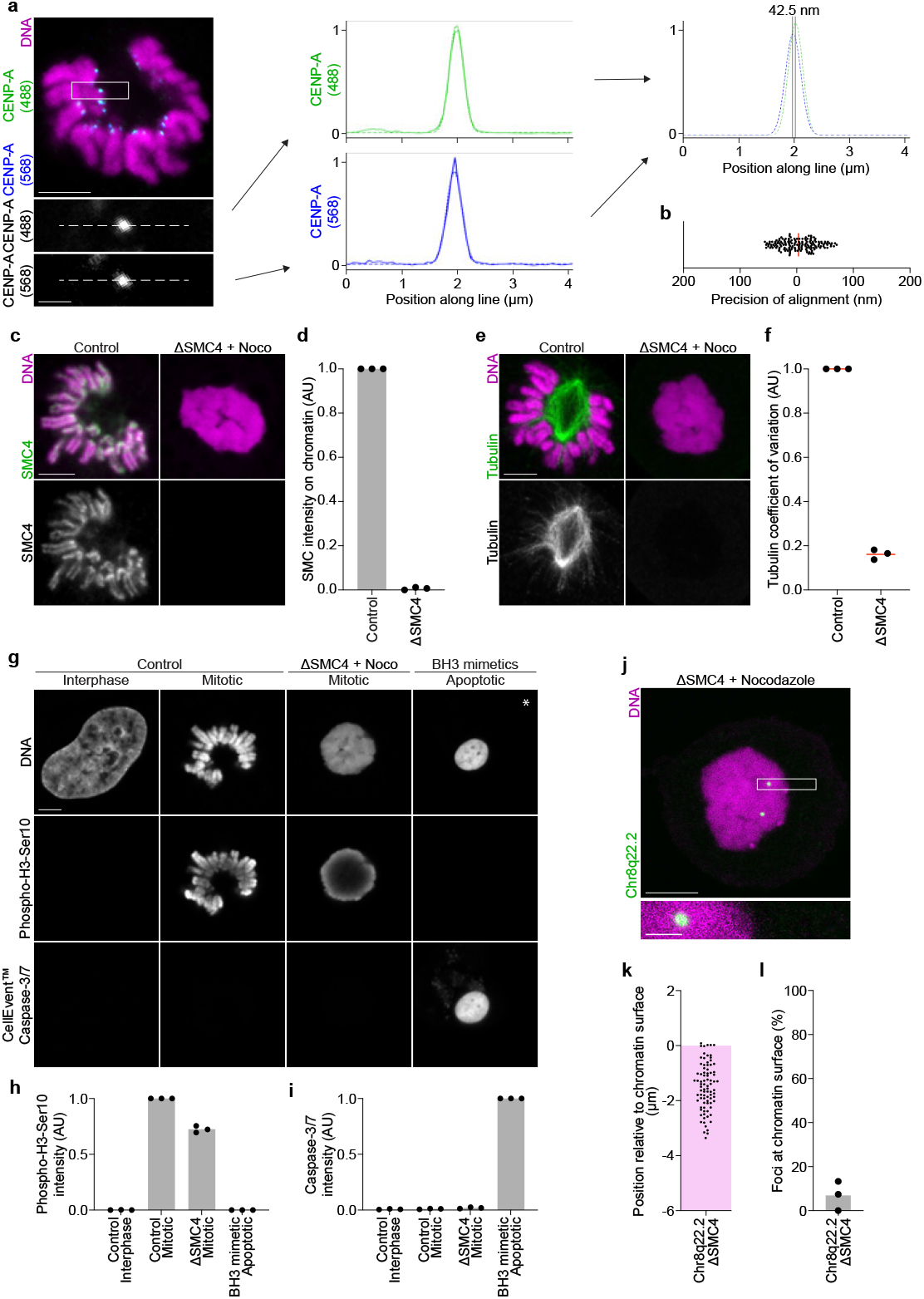
Validation of localisation quantification, condensin-depletion, and spindle depolymerisation. **a**, Overview of linescan analysis procedure. **b**, Localisation precision analysis of CENP-A focus detection, based on two fluorophores staining the same CENP-A loci. **c**, Localisation of SMC4 in mitotic SMC4-AID cells, untreated (control) or treated with 5-Ph-IAA (ΔSMC4) and Nocodazole (Noco). SMC4 was stained by HaloTag ligand. **d**, Quantification of condensin signal on chromatin in cells as in **c** by biological replicate. *n* = 5 cells per condition per replicate. **e**, Immunofluorescent staining of Tubulin in control and in ΔSMC4 cells; DNA stained with Hoechst-33342. **f**, Quantification of microtubule depolymerisation based on Tubulin coefficient of variation from cells treated as in **e** by biological replicate (*n* = 5 / 5 / 4 cells for control and ΔSMC4). Bar indicates mean. **g**, Validating live mitotic state of condensin-depleted cells with depolymerised microtubules. Staining of control (interphase and mitotic) cells, ΔSMC4 cells and cells treated with BH3 mimetics to induce apoptosis as positive controls, using phospho-H3-Ser10 immunofluorescence for mitotic and cleaved-Caspase-3/7 signal by CellEvent^TM^ Green for apoptotic state. DNA visualised by Hoechst-33342 staining. Images are linearly contrast adjusted and matched across conditions, with exception of apoptotic DNA (contrast is three-fold lower to compensate for the high compaction of apoptotic chromatin). **h**, Quantification of phospho-H3-Ser10 immunofluorescent staining as in **g** to assess for mitotic state. **i**, Quantification of cleaved-Caspase-3/7 by CellEvent^TM^ Green signal as in **g** to determine apoptotic state. *n* = 5 cells per condition per replicate. **j**, Localisation of Chr8q22.2 arm locus on ΔSMC4 mitotic chromatin mass, visualised by RASER-FISH and DAPI (DNA). **k**, Quantification of Chr8q22.2 foci localisation relative to chromatin surface in cells treated as described in **j** from three biological replicate (*n* = 30 / 28 / 27 foci from 10 cells each). **l**, Quantification of percentage of foci localised at the chromatin surface as in **j** across biological replicates. Scale bars: 5 µm (**a, c, e, g, j**), 1µm for inset (**a, j**).

**Fig. S2.**
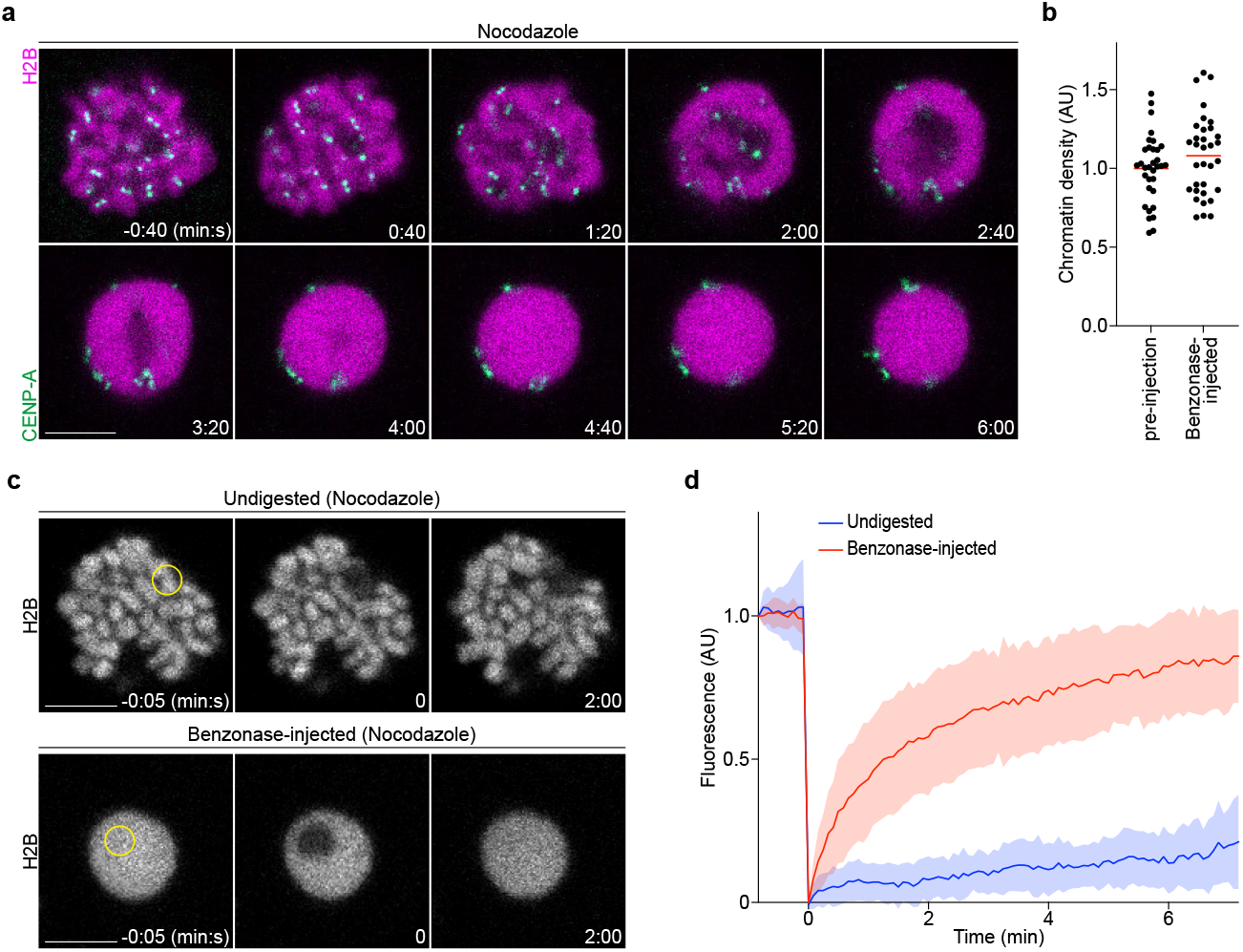
Benzonase-injection results in liquid chromatin condensate. **a**, Chromosome fragmentation in live Nocodazole-arrested mitotic HeLa cells (*t* = 0 min) by Benzonase injection. Central z-section with centromeres visualised by CENP-A-EGFP. Chromatin was visualised by H2B-mCherry. Time is shown as min:s. **b**, Quantification of chromatin density for cells as in **a**. *n* = 11 cells, 3 regions of interest (ROIs) each. Bar indicates mean. **c**, Chromatin mobility in Nocodazole-arrested cells with undigested chromosomes and after Benzonase injection, measured by fluorescence recovery after photobleaching of H2B-mCherry. The circles indicate the photobleaching region at *t* = 0 s. Time is shown as min:s. **d**, Quantification of fluorescence in *n* = 15 undigested and Benzonase-digested cells each across three biological replicates as described in **c**. Data are mean ± s.d.

**Fig. S3.**
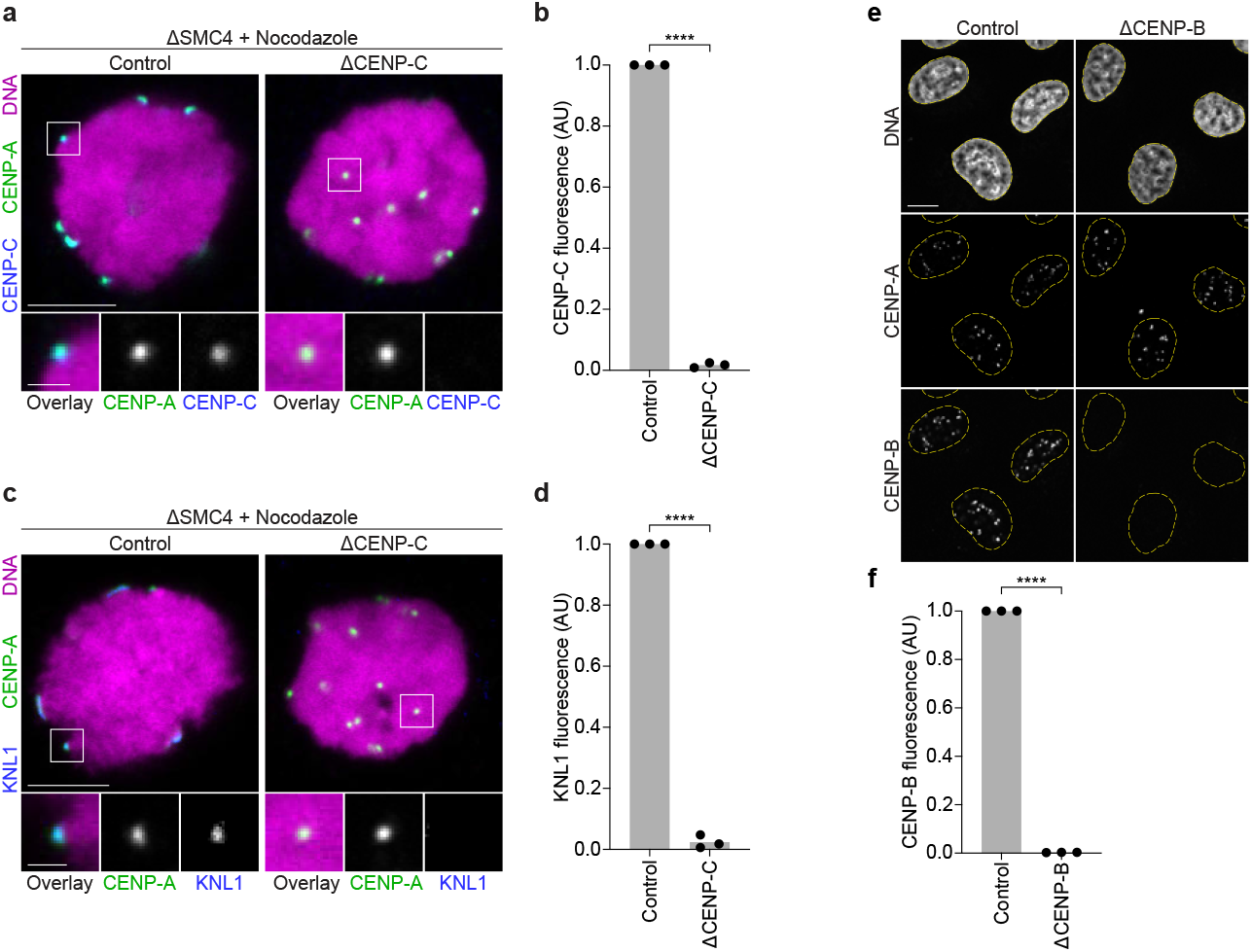
Characterisation of kinetochore depletion and CENP-B knockout. **a**, Immunofluorescence staining for CENP-A and CENP-C after kinetochore-depletion in condensin-depleted (ΔSMC4), microtubule depolymerised cells arrested in mitosis by proTAME; Chromatin was visualised by Hoechst-33342. Scale bars: 5µm, 1µm for inset. **b**, Quantification of mean CENP-C fluorescence on CENP-A in control and ΔCENP-C cells per biological replicate (*n* = 3). Each biological replicate includes 10 foci in 5 cells per condition. **c**, Immunofluorescence staining for CENP-A and KNL1 in control and ΔCENP-C ΔSMC4 cells as in **a**; Chromatin was visualised by Hoechst-33342. Scale bars: 5µm, 1µm for inset. **d**, Quantification of mean KNL1 fluorescence on CENP-A in control and ΔCENP-C cells per biological replicate (*n* = 3). Each biological replicate includes 10 foci in 5 cells per condition. **e**, Immunofluorescence staining for CENP-A and CENP-B in control and ΔCENP-B cell line. Scale bar: 10µm. **f**, Quantification of mean CENP-B fluorescence on CENP-A in control and ΔCENP-B cells per biological replicate (*n* = 3). Each biological replicate includes *n* = 5 / 5 / 5 (control) and *n* = 5 / 5 / 2 (ΔCENP-B) fields of view of ∼15 cells. Significance was tested in all conditions using a two-tailed t-test (**b**: *P* < 10^-15^ (precision limit of floating-point arithmetic), t(4) = 210.5; **d**: *P* < 10^-15^ (precision limit of floating-point arithmetic), t(4) = 78.9; **f**: *P* < 10^-15^ (precision limit of floating-point arithmetic), t(4) = 892.6).

**Fig. S4.**
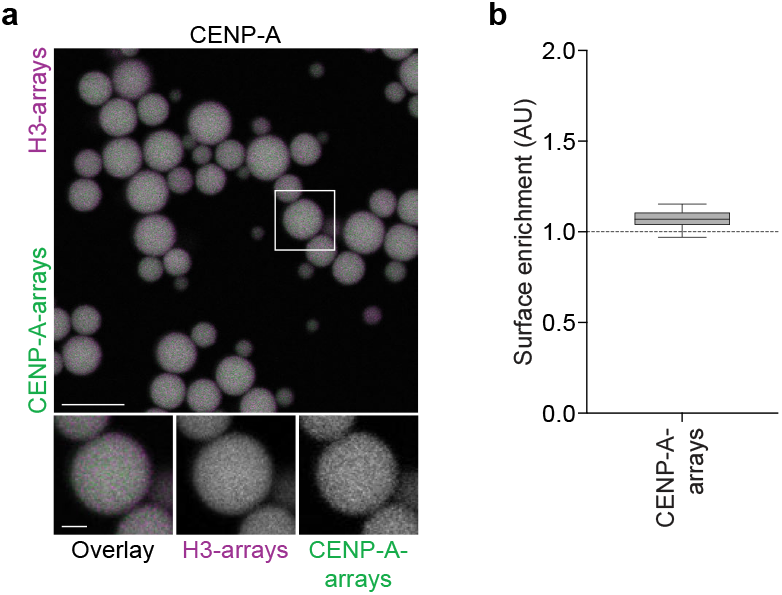
CENP-A alone is not sufficient for surface localisation *in vitro*. **a**, Chromatin condensates of canonical H3-containing nucleosome arrays mixed at 1:1 ratio with CENP-A-containing arrays. H3-arrays visualised by Alexa Fluor 594, CENP-A-arrays visualised by Alexa Fluor 488. Scale bars: 10 µm, 2 µm for inset. **b**, Quantification of CENP-A-arrays surface enrichment for condensates as in **a**. *n* = 296; *n* indicates number of condensates.

**Fig. S5.**
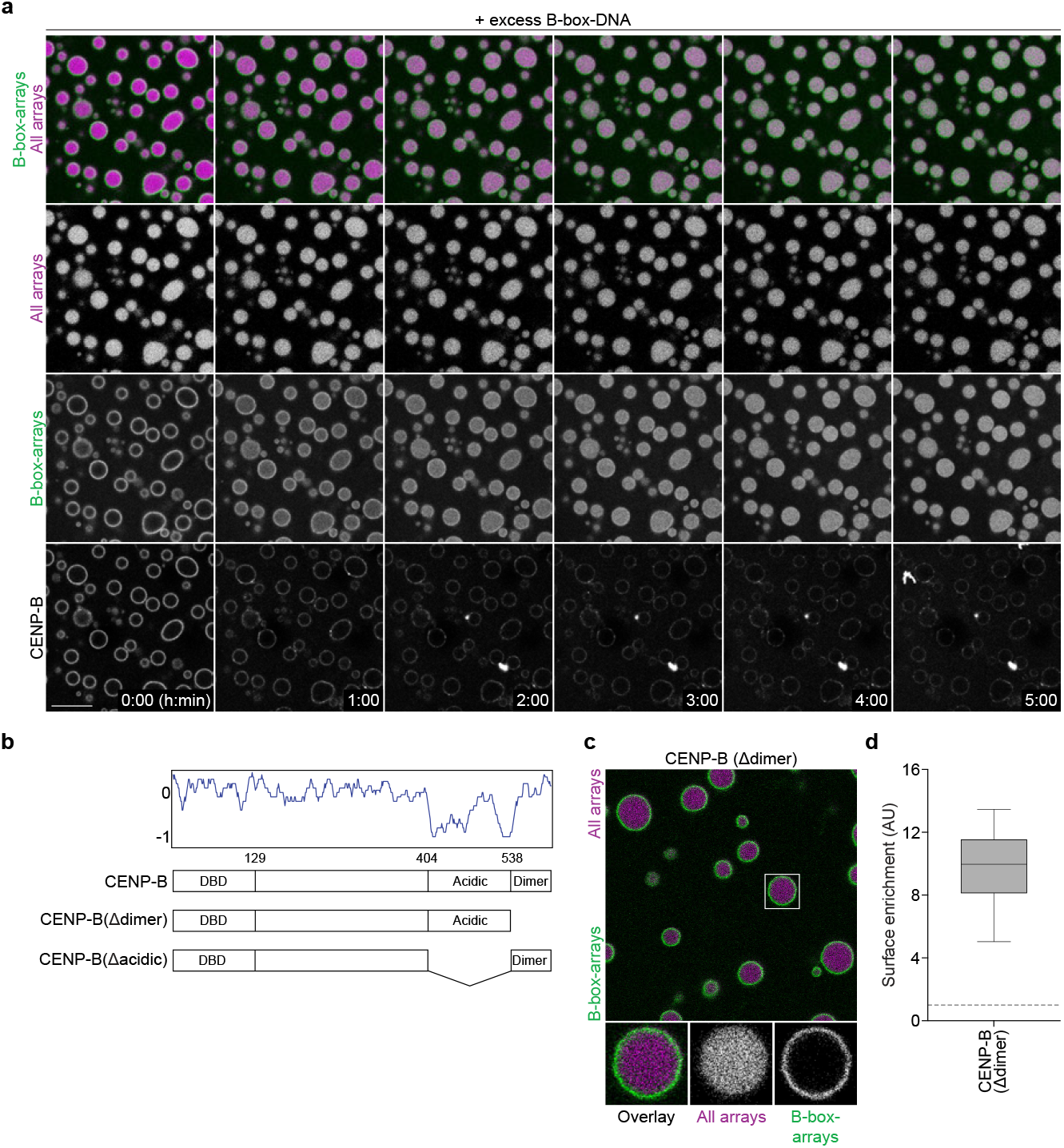
Surface targeting of B-box-nucleosome array depends on CENP-B binding, but not on CENP-B’s dimerisation domain. **a**, Addition of excess B-box-arrays to experiment as in **Fig. 3a**. All arrays were visualised by Hoechst-33342, B-box-arrays by 5’ Cy5, CENP-B by Alexa Fluor 488. Time is shown as h:min. Scale bar: 10µm. **b**, Electrostatic charge profile of CENP-B with the structure of full-length CENP-B, CENP-B lacking the dimerisation domain (Δdimer) or CENP-B lacking the acidic region (Δacidic). **c**, Chromatin condensates of 91% canonical nucleosome arrays and 9% B-box-arrays with full-length CENP-B or with CENP-B lacking the dimerisation domain (Δdimer). All arrays stained with Hoechst-33342, B-box-arrays additionally visualised by 5’ Cy5. Scale bars: 10µm, 2µm for inset. **d**, Quantification of B-box-arrays surface enrichment for condensates as in **c**. *n* = 303 (CENP-B (Δdimer)); *n* indicates number of condensate.

**Fig. S6.**
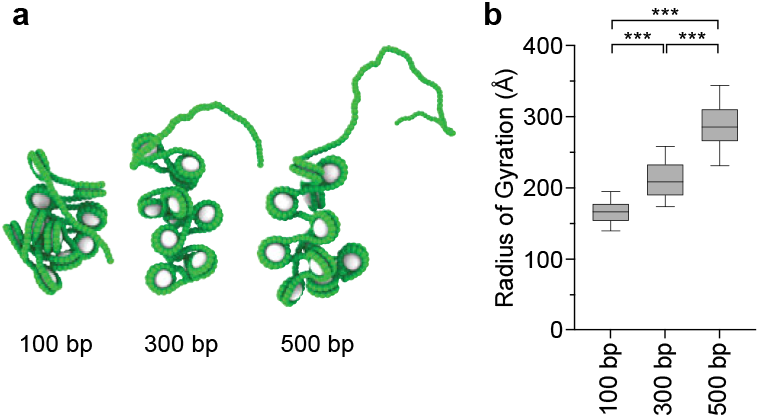
Conformation and energetic analysis of nucleosome arrays with extended DNA segments. **a**, Single molecule representative snapshots of the coarse-grained nucleosome arrays fused to nucleosome-free DNA segments of varying lengths (DNA-arrays). **b**, Distribution of the radius of gyration for DNA-arrays of varying lengths as in **a**. See methods for details on *n*. Significance was tested using two-tailed t-tests (100 bp vs 300 bp: *P* < 10^-15^ (precision limit of floating-point arithmetic), t(1002) = 32; 300 bp vs 500 bp: *P* < 10^-15^ (precision limit of floating-point arithmetic), t(1002) = 39.2; 100 bp vs 500 bp: *P* < 10^-15^ (precision limit of floating-point arithmetic), t(1002) = 73).

**Fig. S7.**
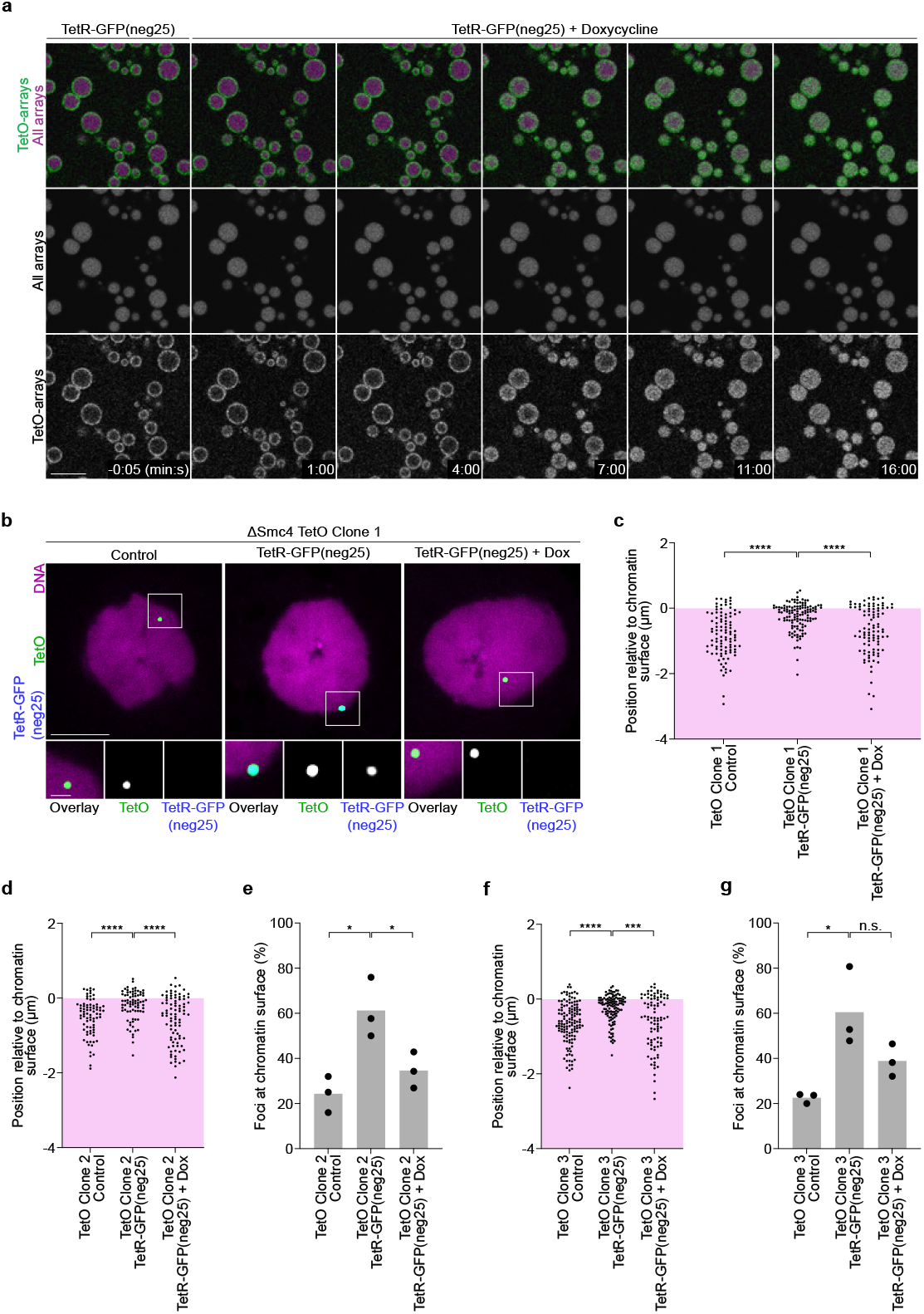
Further characterisation of chromatin surface targeting by TetR-GFP(neg25) *in vitro* and in cells. **a**, Localisation of TetO-arrays when bound by TetR-GFP(neg25), before and after addition of Doxycycline (Dox). All arrays were visualised by Hoechst-33342, TetO-arrays by xx. Time is shown as min:s. Scale bar: 10µm. **b**, Localisation of TetO repeats on ΔSMC4 mitotic chromatin mass for TetO Clone 1, visualised by RASER-FISH (TetO), immunofluorescence (TetR-GFP(neg25)) and DAPI (DNA) in the absence of protein (control), bound by TetR-GFP(neg25), or in the presence of unbound TetR-GFP(neg25) (+ Dox). Scale bars: 5µm, 1µm for inset. **c**, Quantification of TetO localisation relative to chromatin surface for cells as in **a** from three biological replicates. *n* = 98 (control), *n* = 122 (TetR-GFP(neg25)), *n* = 98 (TetR-GFP(neg25) + Dox). Significance was tested using a two-tailed Mann-Whitney *U*-test (Control vs TetR-GFP(neg25): *P* = 6.93 x 10^-11^, U = 2917; TetR-GFP(neg25) vs TetR-GFP(neg25) + Dox: *P* = 6.36 x 10^-11^, U = 4101). **d**, As in **c** for TetO Clone 2. *n* = 78 (control), *n* = 77 (TetR-GFP(neg25)), *n* = 89 (TetR-GFP(neg25) + Dox). Significance was tested using a two-tailed Mann-Whitney *U*-test (Control vs TetR-GFP(neg25): *P* = 1.06 x 10^-5^, U = 1772; TetR-GFP(neg25) vs TetR-GFP(neg25) + Dox: *P* = 8.69 x 10^-5^, U = 2214). **e** Quantification of percentage of foci localised at the chromatin surface as in **d** per biological replicate (*n* = 3) across conditions. *n* = 25 / 28 / 25 (control), *n* = 26 / 26 / 25 (TetR-GFP(neg25)), *n* = 35 / 28 / 26 (TetR-GFP(neg25) + Dox), with *n* indicating number of foci/cells per biological replicate. Significance was tested using a two-tailed t-test (Control vs TetR-GFP(neg25): *P* = 0.0148, t(4) = 4.1; TetR-GFP(neg25) vs TetR-GFP(neg25) + Dox: *P* = 0.0418, t(4) = 3.0). **f**, As in **c** for TetO Clone 3. *n* = 131 (control), *n* = 119 (TetR-GFP(neg25)), *n* = 90 (TetR-GFP(neg25) + Dox). Significance was tested using a two-tailed Mann-Whitney *U*-test (Control vs TetR-GFP(neg25): *P* = 6.33 x 10^-10^, U = 4264; TetR-GFP(neg25) vs TetR-GFP(neg25) + Dox: *P* = 4.72 x 10^-4^, U = 3841). **g**, Quantification of percentage of foci localised at the chromatin surface as in **f** per biological replicate (*n* = 3) across conditions. *n* = 25 / 76 / 30 (control), *n* = 26 / 70 / 23 (TetR-GFP(neg25)), *n* = 34 / 28 / 28 (TetR-GFP(neg25) + Dox), with *n* indicating number of foci/cells per biological replicate. Significance was tested using a two-tailed t-test (Control vs TetR-GFP(neg25): *P* = 0.0213, t(4) = 3.7; TetR-GFP(neg25) vs TetR-GFP(neg25) + Dox: *P* = 0.1229, t(4) = 1.95).

**Table S1.** Charge of kinetochore proteins associated per single CCAN at physiological pH (= 7.4).

| Name | Stoichiometry | pI | charge |
| --- | --- | --- | --- |
| CENP-H | 1 | 5.23 | -7.1 |
| CENP-I | 1 | 8.98 | 18.3 |
| CENP-K | 1 | 4.83 | -18.8 |
| CENP-M | 1 | 6.7 | -0.7 |
| CENP-N | 1 | 9.18 | 8.5 |
| CENP-L | 1 | 6.07 | -4.8 |
| CENP-C | 1 | 9.43 | 36.7 |
| CENP-T | 1 | 6.14 | -6.7 |
| CENP-W | 1 | 11.29 | 17.9 |
| CENP-S | 1 | 5.83 | -2.2 |
| CENP-X | 1 | 5.59 | -2 |
| CENP-O | 1 | 7.63 | 1.3 |
| CENP-P | 1 | 5.9 | -4.8 |
| CENP-Q | 1 | 9.43 | 10.4 |
| CENP-U | 1 | 9.18 | 11.7 |
| CENP-R | 1 | 9.13 | 5.2 |
| KNL1 | 2 | 5.3 | -81.4 |
| Zwint | 2 | 5.1 | -10 |
| DSN1 | 2 | 6.57 | -1.6 |
| MIS12 | 2 | 5.5 | -5.2 |
| NSL1 | 2 | 4.68 | -17.6 |
| PMF1 | 2 | 5.37 | -6 |
| NDC80 | 4 | 5.48 | -17.5 |
| NUF2 | 4 | 8.41 | 4.4 |
| SPC24 | 4 | 4.65 | -18.6 |
| SPC25 | 4 | 7.7 | 0.9 |
| **Total** |  |  | **-303.9** |

